# A nutrient-dependent division antagonist is regulated post-translationally by the Clp proteases in *Bacillus subtilis*

**DOI:** 10.1101/176222

**Authors:** Norbert S. Hill, Jason D. Zuke, P. J. Buske, An-Chun Chien, Petra Anne Levin

## Abstract

Changes in nutrient availability have dramatic and well-defined impacts on both transcription and translation in bacterial cells. At the same time, the role of post-translational control in adaptation to nutrient-poor environments is poorly understood. Here we report a role for the bacterial Clp proteases in degradation of the division inhibitor UgtP during growth in nutrient-poor medium. Under nutrient-rich conditions, interactions with its substrate UDP-glucose promote interactions between UgtP and the tubulin-like cell division protein FtsZ in *Bacillus subtilis*, inhibiting maturation of the cytokinetic ring and increasing cell size. In nutrient-poor medium, reductions in UDP-glucose availability favor UgtP oligomerization, sequestering it from FtsZ and allowing division to occur at a smaller cell mass. Intriguingly, in nutrient-poor conditions UgtP levels are reduced ∼ 3-fold independent of UDP-glucose, suggesting an additional layer of regulation. *B. subtilis* cells cultured under different nutrient conditions indicate that UgtP accumulation is controlled through a nutrient-dependent post-translational mechanism dependent on the Clp proteases. Notably, all three *B. subtilis* Clp chaperones appeared able to target UgtP for degradation during growth in nutrient-poor conditions. Together these findings highlight conditional proteolysis as a mechanism for bacterial adaptation to a rapidly changing nutritional landscape.

## BACKGROUND

As single-celled organisms, bacteria must constantly alter their physiology to adapt to their environment. Nutrients in particular can dramatically impact bacterial growth and morphology. *Escherichia coli*, *Salmonella*, and *Bacillus subtilis* cells grow several times faster and are up to three times larger when cultured in nutrient-rich medium than when cultured in nutrient-poor medium (1-3). Nutrient-dependent increases in cell size appear to a means of accommodating the concomitant increase in macromolecular biosynthesis at faster growth rates, particularly the additional DNA generated by multifork replication (4, 5).

The nutrient-dependent regulation of biosynthesis has been an area of intense interest for many years (6). Numerous studies have explored how changes in nutrient composition and growth rate impact transcription and translation, which in large part is a response mediated via accumulation of the signaling molecule guanosine pentaphosphate ((p)ppGpp) (7-12). Although post-translational regulation has been implicated in adaptation to changes in growth phase (e.g. carbon starvation (13-15)), how fluctuations in nutritional content and growth rate impact post-translational regulation at the molecular level is poorly defined.

In previous work, we identified a class of division antagonists responsible for coordinating cell size with nutrient availability in *B. subtilis* and *E. coli* (4, 5). Both organisms employ unrelated, yet functionally similar, glucosyltransferases—UgtP in *B. subtilis* and OpgH in *E. coli*—to inhibit division and increase size during growth in carbon-rich medium (16, 17). In both cases, binding to their substrate, UDP-glucose, stimulates interaction between UgtP and OpgH and the tubulin-like cell division protein FtsZ to delay the maturation of the cytokinetic ring and increase cell size. Loss-of-function mutations in *ugtP* or *opgH* and in genes required for UDP-glucose biosynthesis reduce cell size by as much as 35% during growth in nutrient-rich conditions.

UgtP and OpgH are both moonlighting proteins with additional roles as glucosyltransferases that contribute to cell envelope biogenesis. UgtP transfers glucose moieties from UDP-glucose to diacylglycerol to form the diglucosyl-diacylglycerol membrane anchor for lipoteichoic acid (LTA) (18). OpgH also binds the nucleotide sugar UDP-glucose to transfer glucose into the periplasm as an initial step toward the synthesis of osmoregulated periplasmic glucans (OPGs) (19). LTA and OPGs are proposed to have similar functions (20) based on the conservation of enzymes involved in their synthesis, their location within the cellular envelope (21, 22), and their potential contribution to osmoprotection (17, 23).

In *B. subtilis*, UDP-glucose increases UgtP’s affinity for FtsZ (24). During growth in nutrient-rich conditions UgtP is localized throughout the cytoplasm, where the largest pool of FtsZ is located, and can also be found at the cytokinetic ring and at cell poles (4). During growth in carbon-poor conditions or when synthesis of UDP-glucose is disrupted, UgtP self-assembles into large oligomers, sequestering it from FtsZ and permitting division to occur at a reduced cell size. *In vitro* studies suggest UDP-glucose acts as a molecular rheostat, precisely modulating UgtP’s affinity for itself and FtsZ to coordinate size with growth rate and nutrient availability (24). Curiously, while the UgtP homolog in the Gram-positive pathogen *Staphylococcus aureus* interacts with FtsZ and other divisome proteins, it does not exhibit the same dynamic localization pattern it does in *B. subtilis* nor does it appear to make a significant contribution to cell size (21).

In addition to UDP-glucose-dependent changes in its affinity for FtsZ, UgtP is also subject to nutrient-dependent changes in concentration. UgtP levels are reduced several-fold during growth in nutrient-poor conditions (4). Defects in the UDP-glucose biosynthesis pathway have no discernable impact on the intracellular concentration of UgtP, suggesting that nutrient-dependent changes in accumulation are independent of the signaling molecule (4).

The striking difference in UgtP levels, together with previous work suggesting protein turnover might be increased in nutrient-poor conditions (25), prompted us to investigate the mechanism underlying this additional layer of UgtP regulation. Here we report that UgtP nutrient-dependent accumulation is governed by a post-translational mechanism involving all three substrate recognition components of the *B. subtilis* Clp protease system. We find that the *clp* genes themselves are upregulated during growth in nutrient-poor medium, suggesting a possible mechanism for increased UgtP degradation under these conditions. These findings suggest an important role for conditional proteolysis in the nutrient-dependent regulation of cellular processes.

## RESULTS

### UgtP accumulation is subject to nutrient-dependent post-translational regulation

In our initial investigation, we observed that the intracellular concentration of a UgtP-6XHis fusion protein was several times lower when cells were cultured under nutrient-poor conditions, but unaffected by the absence of UDP-glucose (4). To expand on this observation we measured levels of the same UgtP-His fusion across four different nutrient conditions: Lysogeny Broth (LB), S7_50_ + 1% glucose (minimal glucose), S7_50_ + 1% glycerol (minimal glycerol), and S7_50_ + 1% sorbitol (minimal sorbitol). Under each condition, the respective mass doubling time of this strain (*P*_*ugtP*_*-ugtP-his*) was 22’, 39’, 58’, and 78’.

Consistent with our previous findings, UgtP-His levels increased linearly with growth rate, as evidenced by a quantitative immunoblot probed with an α-His antibody (Fig. 1). The intracellular concentration of UgtP-His was ∼ 3-fold lower in cells cultured in minimal sorbitol, the most nutrient-poor condition, than in those cultured in LB, the most nutrient-rich condition. To control for the possibility that the His-tag was impacting the stability of UgtP-His, we also measured YFP-UgtP levels in a strain expressing the fusion protein from a xylose-inducible promoter (*P*_*xyl*_*-yfp-ugtP*), cultured in both LB + 0.5% xylose and minimal sorbitol + 0.5% xylose. As we observed with UgtP-His, YFP-UgtP levels were ∼3-fold lower in minimal sorbitol compared to LB supporting a model in which UgtP (and not the His or YFP tag) is the primary target for degradation during growth in nutrient-poor medium (Additional File 1: Fig. S1).

**Figure 1.**
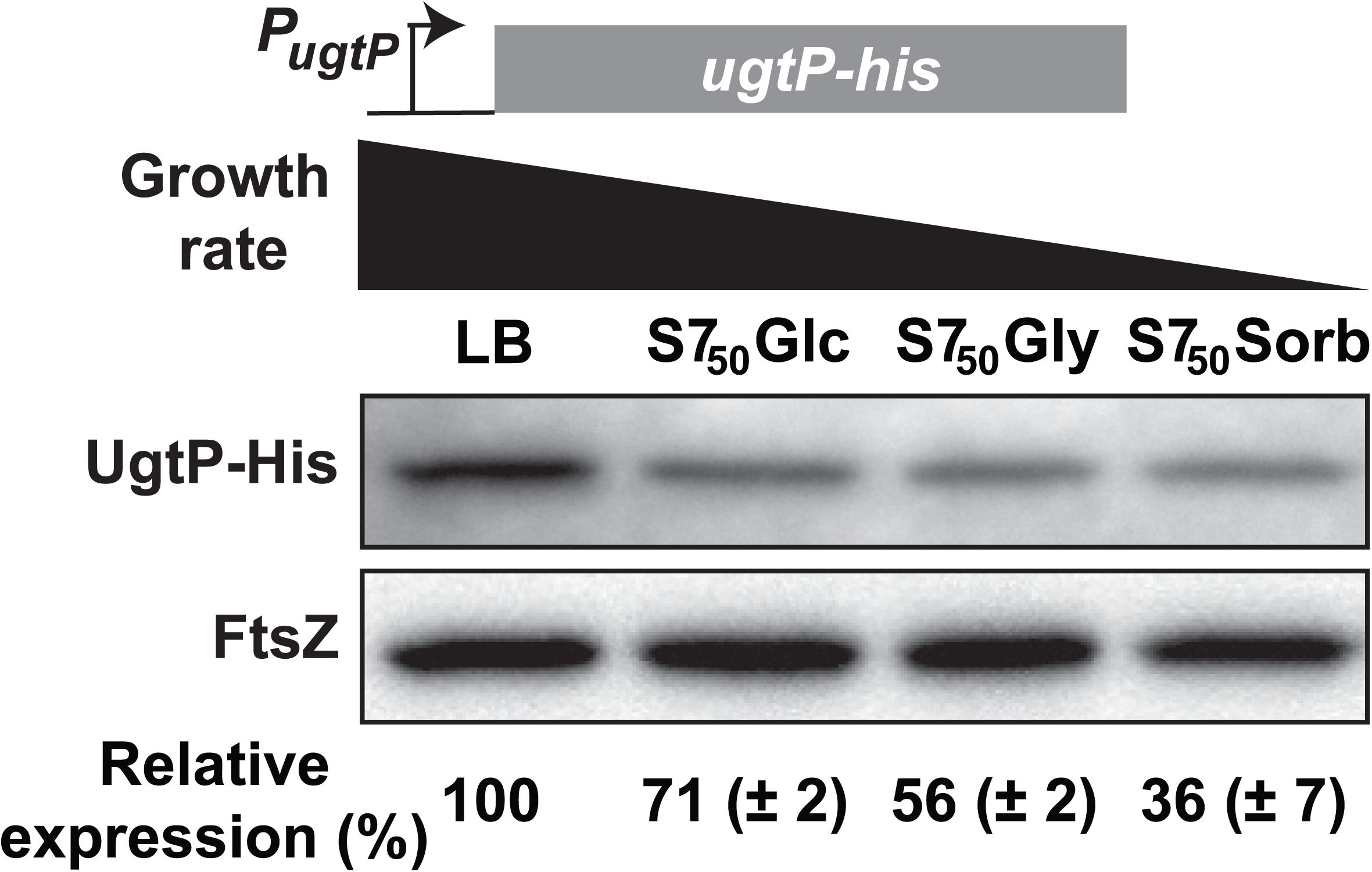
UgtP accumulates only in nutrient-rich conditions. Quantitative immunoblot measuring native UgtP-His levels in a range of growth conditions. PL2265 (*P*_*ugtP*_*-ugtP-his*) was cultured in LB (τ = 22’), minimal glucose (τ = 39’), minimal glycerol (τ = 58’), or minimal sorbitol (τ = 78’). FtsZ is shown as a loading control. Protein levels of LB are set as the reference in the relative expression below (n = 3, error = SD).

To determine if UgtP is subject to transcriptional or post-transcriptional modes of regulation, we generated two *ugtP-lacZ* fusion constructs. In the first construct, a reporter for *ugtP* transcription, the 700 base pairs immediately upstream of the *ugtP* start codon were fused to *lacZ*, leaving the *lacZ* Shine-Dalgarno sequence intact. In the second construct, a reporter for UgtP translation, *lacZ* was fused in-frame downstream of the first 30 codons of the *ugtP* open reading frame that included the native *ugtP* Shine-Dalgarno sequence.

*lacZ* expression data suggest that nutrient-dependent changes in the intracellular level of UgtP are independent of both transcriptional and translational control. In stark opposition to UgtP-His levels, expression of both the transcriptional and translational *lacZ* fusions was inversely proportional to growth rate. *lacZ* expression from the transcriptional fusion was 2-fold higher in cells cultured in minimal sorbitol than LB (Fig. 2A) and expression from the translational fusion was 4-fold higher in minimal sorbitol than LB (Fig. 2B). qRT-PCR data also indicated that *ugtP* expression is elevated during growth in nutrient-poor conditions (Fig. 2C). *ugtP* levels were 1.3-fold higher in minimal glucose, 1.7-fold higher in minimal glycerol, and 2.5-fold higher in minimal sorbitol compared to LB. In all, these data strongly support a model where nutrient-dependent changes in UgtP accumulation are governed by a post-translational mechanism.

**Figure 2.**
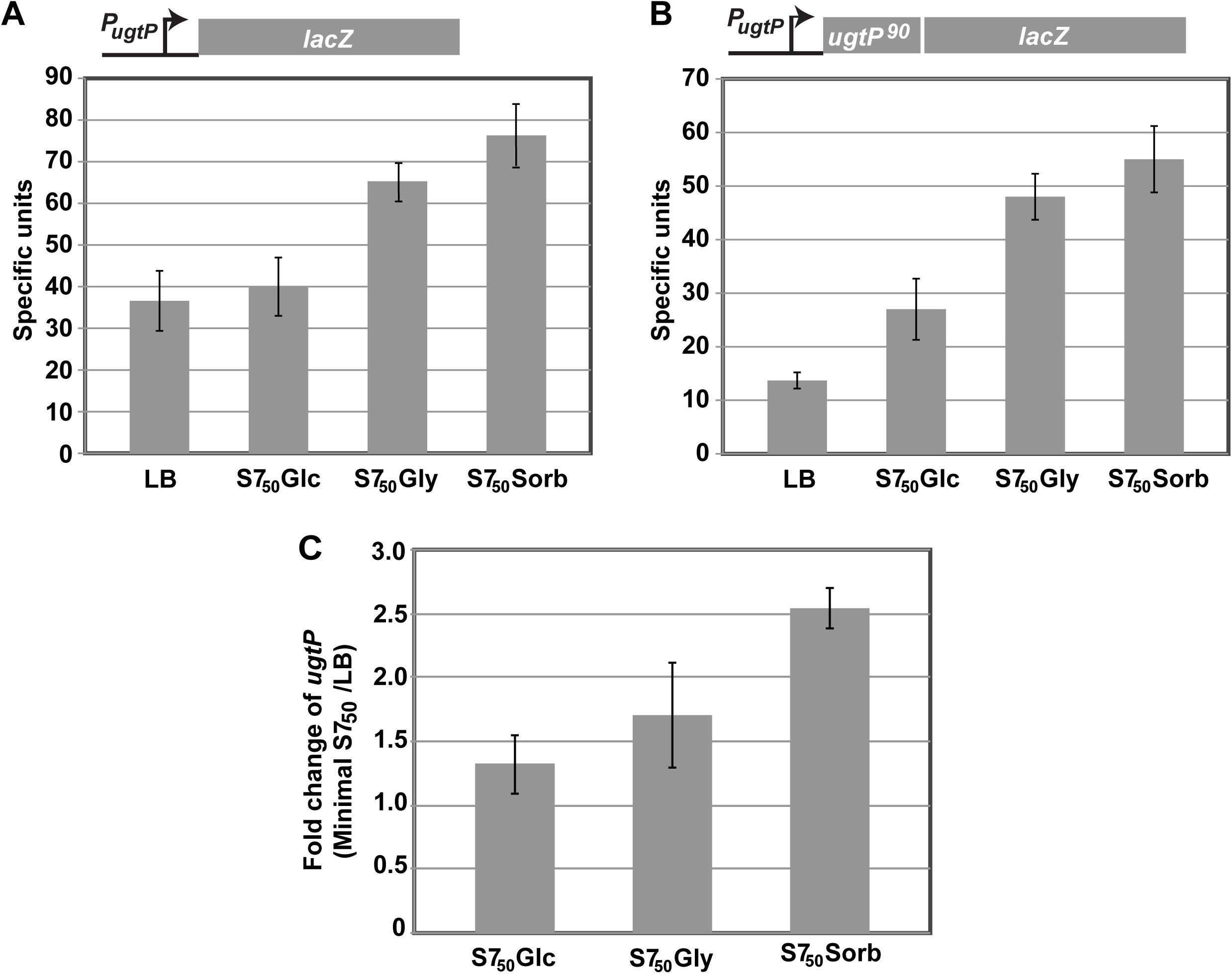
UgtP accumulation is subject to post-translation control. *lacZ* encoding β-galactosidase is fused either **(A)** 700bp upstream of the *ugtP* start site (*P*_*ugtP*_) to generate a transcriptional fusion (PL1967) or **(B)** an additional 90 bases downstream of the ugtP start codon to generate a translational fusion (PL2034). Both strains were cultured in a range of nutrient conditions to generate four different growth rates. Bars indicate specific β-galactosidase activity (n = 3, error = SD). **(C)** qRT-PCR measurements of *ugtP* expression levels. Expression in three defined media is normalized to expression in LB (n = 3, error = SD).

### The Clp proteases are responsible for UgtP degradation under nutrient-poor conditions

To understand the mechanism responsible for the post-translational regulation of UgtP, we sought to identify the protease responsible for its degradation. As an initial step, we screened *B. subtilis* strains defective in five well-studied proteases: YluC, CptA, ClpP, YvjB, and Lon for aberrant UgtP regulation. We rationalized that if one of these proteases were responsible for the growth rate-dependent degradation of UgtP, then UgtP would inappropriately accumulate in its absence during growth in nutrient-poor conditions. For this experiment we used the *P*_*xyl*_*-ugtP-his* construct described above to enhance our ability to detect UgtP by quantitative immunoblot.

Analysis of the protease-deficient strains strongly implicated ClpP in the nutrient-dependent control of UgtP accumulation. UgtP-His levels increased significantly under nutrient-poor conditions only in the strain defective for ClpP (Fig. 3A). UgtP-His levels were approximately 5-fold higher during growth in minimal sorbitol in the Δ*clpP* strain than in the parental strain or in the four other protease mutants.

**Figure 3.**
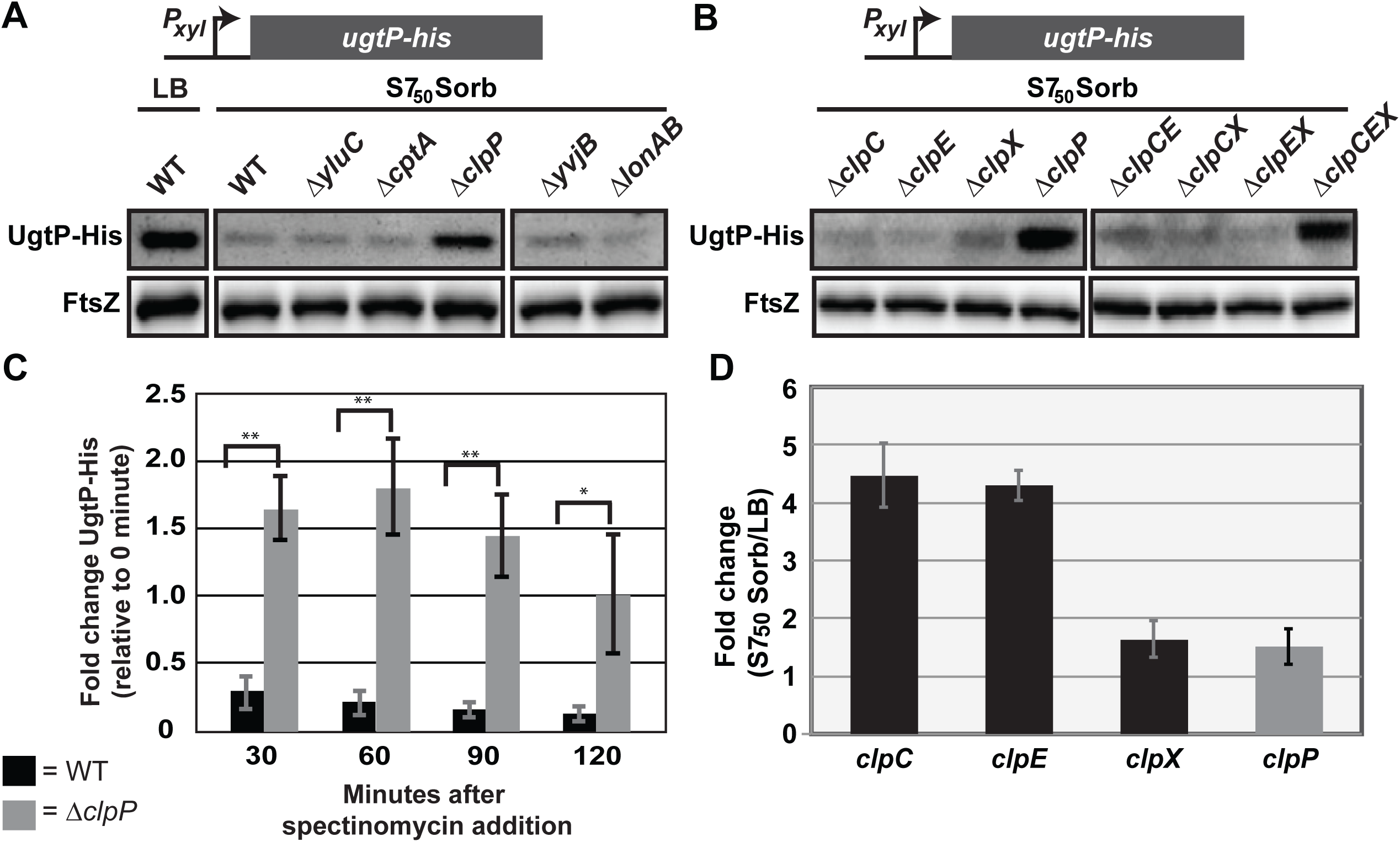
UgtP is subject to nutrient-dependent, post-translational regulation by the Clp proteases. Quantitative immunoblot of UgtP-His expressed from a xylose-inducible promoter (*P*_*xyl*_*-ugtP-his*) in the absence of **(A)** 5 proteases, YluC, CptA, ClpP, YvjB, and Lon (PL2022, PL2028, PL2102, PL2032, and PL2033, respectively and BH10 serving as WT) or **(B)** single and combinatorial deletions of the three Clp chaperones (ClpC, ClpE, and/or ClpX) (BH127, BH128, BH130, PL2102, BH135, BH136, BH137, BH138). Cells were cultured in minimal sorbitol + 0.5% xylose, and immunoblotted against His and FtsZ (loading control). **(C)** Fold change in UgtP-His levels for BH10 (*P*_*xyl*_*-ugtP-his*) and BH129 (*P*_*xyl*_*-ugtP-his*; *ΔclpP*) after the addition of spectinomycin to inhibit translation. Cells were cultured in minimal sorbitol + 0.5% xylose, sampled every 30 minutes after spectinomycin addition, and subjected to immunoblotting against His antibody. * indicates p < 0.05, ** indicates p < 0.005 (n = 3, error = SD). T-test analysis was used to assess differences in UgtP-His levels between strains. **(D)** qRT-PCR measurements of *clpC*, *clpE*, *clpX*, and *clpP* expression in minimal sorbitol versus LB (n = 3, error = SD).

ClpP is a processive serine protease widely conserved throughout bacteria. Ineffective on its own, ClpP functions in tandem with AAA+ ATPase Clp chaperones (26). The Clp chaperones are responsible for recognizing target proteins, unfolding them using the energy of ATP hydrolysis, and translocating the unfolded polypeptide into the ClpP proteolytic chamber (27). In *B. subtilis*, there are three Clp chaperones: ClpC, ClpE, and ClpX. Since each chaperone identifies a unique set of substrates with limited target overlap, we reasoned that either ClpC, ClpE, or ClpX would be responsible for growth rate-dependent degradation of UgtP. To distinguish the Clp chaperone involved, we examined accumulation of UgtP-His from a xylose-inducible promoter in Δ*clpC*, Δ*clpE*, and Δ*clpX* cells cultured in nutrient-poor medium.

Deletion of *clpC*, *clpE*, and *clpX* alone or in pairwise combination had little impact on UgtP-His levels in minimal sorbitol. UgtP-His accumulated to levels similar to that observed in the Δ*clpP* mutant only in a strain defective for all three chaperones (Fig. 3B). These data suggest that all three Clp chaperones function redundantly to control UgtP accumulation in a growth rate-dependent manner. Multiple proteases targeting a single client protein is reminiscent of the posttranslational regulation of ZapC, a positive regulator of cell division in *E. coli* whose stability is dependent on both ClpXP and ClpAP (28)

To confirm that the Clp proteases are indeed responsible for degradation of UgtP, as our data suggested, we employed an *in vivo* proteolysis assay similar to that described in (29). Briefly, we cultured *P*_*xyl*_*-ugtP-his* and the congenic *ΔclpP* strains in minimal sorbitol with xylose, inhibited new protein synthesis once mid-log was reached, sampled cells every 30 minutes for 2 hours, and then measured UgtP-His levels via quantitative immunoblot. Consistent with UgtP being directly targeted by ClpP, UgtP-His levels decreased rapidly in the presence of ClpP over the time course, but were stable in the absence of ClpP (Fig. 3C and Additional File 2: Fig. S2).

Consistent with the Clp proteases serving as nutrient-dependent regulators of protein stability, expression of all three *clp* chaperone genes and *clpP* were elevated in minimal sorbitol, providing a possible explanation for growth medium-dependent changes in UgtP accumulation (Fig. 3D). qRT-PCR indicated that expression of *clpC* and *clpE* is upregulated ∼ 4-fold, while *clpX* and *clpP* are upregulated ∼1.5 fold in minimal sorbitol, our poorest nutrient condition, relative to LB. These findings are consistent with those of the Hecker lab, who observed ClpXP and ClpCP drive proteolysis of a large group of proteins, most notably enzymes involved in central carbon metabolism, upon glucose starvation (UgtP was not identified as a Clp substrate in this study) (25).

### Defects in UgtP’s hexose-binding site increases susceptibility to proteolysis *in vivo*

In an effort to identify regions of UgtP required for interaction with the Clp chaperones, we screened a series of UgtP mutants for sensitivity to Clp-mediated degradation *in vivo*. Of particular interest were regions of UgtP that mediate interaction with UDP-glucose, FtsZ, or itself. Three putative UgtP mutants were employed for this experiment: one defective in its putative uracil-binding size (ΔURA, F112A V117A), one defective in the putative hexose-binding site (ΔHEX, E306A N309A), and a putative oligomerization mutant (ΔOLI, I142A E146A). Mutations were generated in conserved residues based on structural data from the UgtP homologue, MDGD synthase (30). Structural data indicate that MDGD has two binding sites for UDP-glucose conserved in UgtP: one to coordinate interactions with the uracil moiety on UDP-glucose and one for the hexose moiety. In addition, structural analysis identified a dimerization site on MDGD. We speculated that conserved residues within the analogous region of UgtP are involved in oligomerization. All three UgtP mutants behaved as predicted based on structural and biochemical data from MDGD synthase (30). Both UgtPΔURA and UgtPΔHEX are defective as sugar transferases, exhibit a punctate localization pattern during growth in nutrient-rich medium, and fail to complement a *ugtP* null strain for cell size, all of which is consistent with increased self-association and a loss of interaction with FtsZ (24). In support of our model that oligomerization serves to sequester UgtP from FtsZ under nutrient-poor conditions, the putative oligomerization mutant (UgtPΔOLI) exhibited medial localization (Additional File 3A: **Fig. S3A**) and antagonized division to increase cell size (>30%) in the absence of UDP-glucose (Additional File 3B: **Fig. S3B**).

For analysis of UgtP mutant accumulation, his-tagged versions of all three UgtP variants (ΔURA, ΔHEX and ΔOLI) were expressed from an ectopic locus under the control of a xylose-inducible promoter in a *ugtP* null background. UgtP mutant accumulation under nutrient-rich and nutrient-poor conditions was monitored by quantitative immunoblot in early exponential phase. Both UgtP-His(ΔURA) and UgtP-His(ΔHEX) exhibited migration patterns different from wild type UgtP-His when separated by SDS-PAGE prior to immunoblotting (Fig. 4). This difference may reflect conformational changes in mutant protein structure.

**Figure 4.**
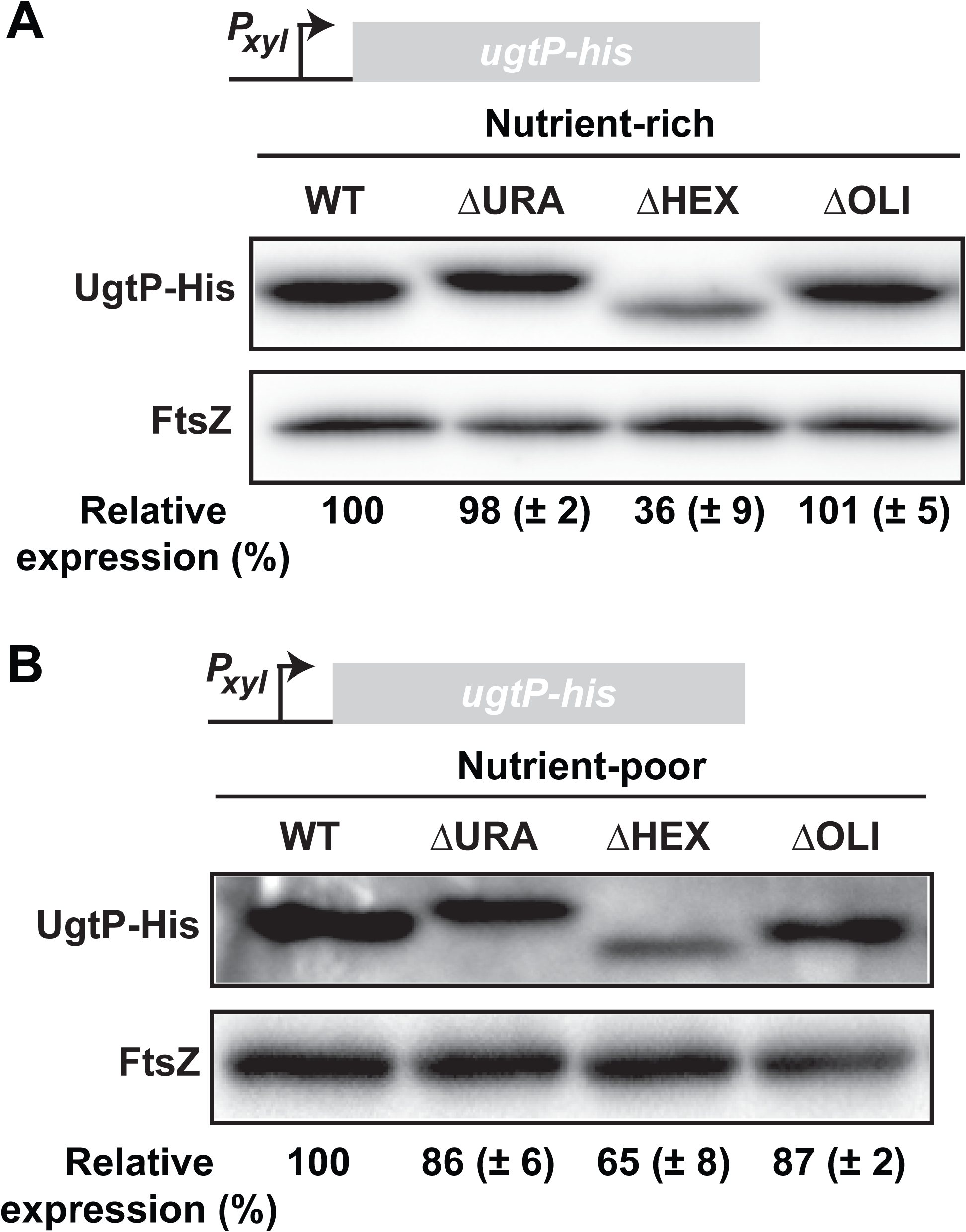
Mutations in UgtP’s putative hexose-binding site enhance susceptibility to proteolysis *in vivo*. UgtP-His variants BH736 (WT), BH742 (ΔURA), BH752 (ΔHEX), BH740 (ΔOLI) were cultured in either **(A)** LB + 0.5% xylose or **(B)** minimal sorbitol + 0.5% xylose and subjected to quantitative immunoblotting against His and FtsZ (loading control) antibodies. Relative expression compared to WT (BH736) UgtP-His is shown (n = 3, error = SD).

Of the three mutants, only UgtP-His(ΔHEX) exhibited ClpP-dependent differential in accumulation compared to wild type UgtP-His, suggesting hexose binding might protect UgtP from proteolysis. In lysate from cells cultured in LB, UgtP-His(ΔHEX) accumulated to only ∼35% of UgtP-His(WT) levels (Fig. 4A). Even in minimal sorbitol UgtP-His(ΔHEX) levels were only ∼65% of UgtP-His(WT) (Fig. 4B). Importantly, each of the constructs exhibited congruent mRNA levels measured by qRT-PCR and UgtP-His(ΔHEX) is at WT levels in a Δ*clpP* background strain in both LB and minimal sorbitol, indicating that the change in stability is directly due to ClpP-mediated degradation (Additional File 4: Fig. S4). In contrast, UgtP-His(ΔURA) and UgtP-His(ΔOLI) did not exhibit ClpP-dependent differential in accumulation, suggesting that neither interaction with FtsZ nor homo-oligomerization impact Clp recognition.

Together these data suggest that interaction with the hexose moiety of UDP-glucose, which we have measured as being more prevalent when cultured in nutrient-rich conditions (Additional File 5: Fig. S5), may provide some protection from Clp-mediated proteolysis. At the same time, however, this observation is inconsistent with our previous observation that UgtP accumulation is independent of UDP-glucose synthesis (4), a point we address in the discussion.

### UgtP is targeted by ClpXP for proteolysis *in vitro*

To clarify whether Clp recognition of UgtP is direct or reliant on a nutrient-dependent adaptor protein, we next determined if UgtP could be degraded directly by ClpXP *in vitro*. Although our genetic data indicate all three Clp chaperones are capable of degrading UgtP (Fig. 3B), we focused on ClpX as the most prevalent chaperone (there are an estimated 1400 ClpX, 250 ClpC, and 100 ClpE hexamers per cell during growth in LB) (31). To ensure our purified ClpXP was functional, we tested whether a well-characterized ClpXP substrate, Spx (32), and a non-targeted protein (Thioredoxin-His) could be proteolyzed. Spx was thoroughly digested in an ATP-dependent manner, while Thioredoxin-His was retractile to ClpXP proteolysis (Additional File 6: Fig. S6). Note that the anti-His antibody interacted strongly with nanogram amounts of the His-UgtP fusion protein (data not shown). Thus, to ensure the experiments were in linear range, we found it necessary to utilize lower concentrations of His-UgtP relative to the Spx and Thioredoxin-His control proteins.

UgtP is degraded by ClpXP *in vitro*, consistent with a direct interaction between this Clp chaperone and UgtP (Fig. 5A). Incubating purified His-UgtP with the ClpXP complex without ATP resulted in minimal degradation (13%). In stark contrast, when ATP was added to reactions 88% of His-UgtP was proteolyzed. Due to the size-similarity between His-UgtP and ClpX used in the degradation assays (44 kDa vs 46 kDa), we could not distinguish between the two when separated by standard SDS-PAGE. Instead, we employed a monoclonal anti-His antibody to distinguish between His-UgtP and ClpX via immunoblot.

**Figure 5.**
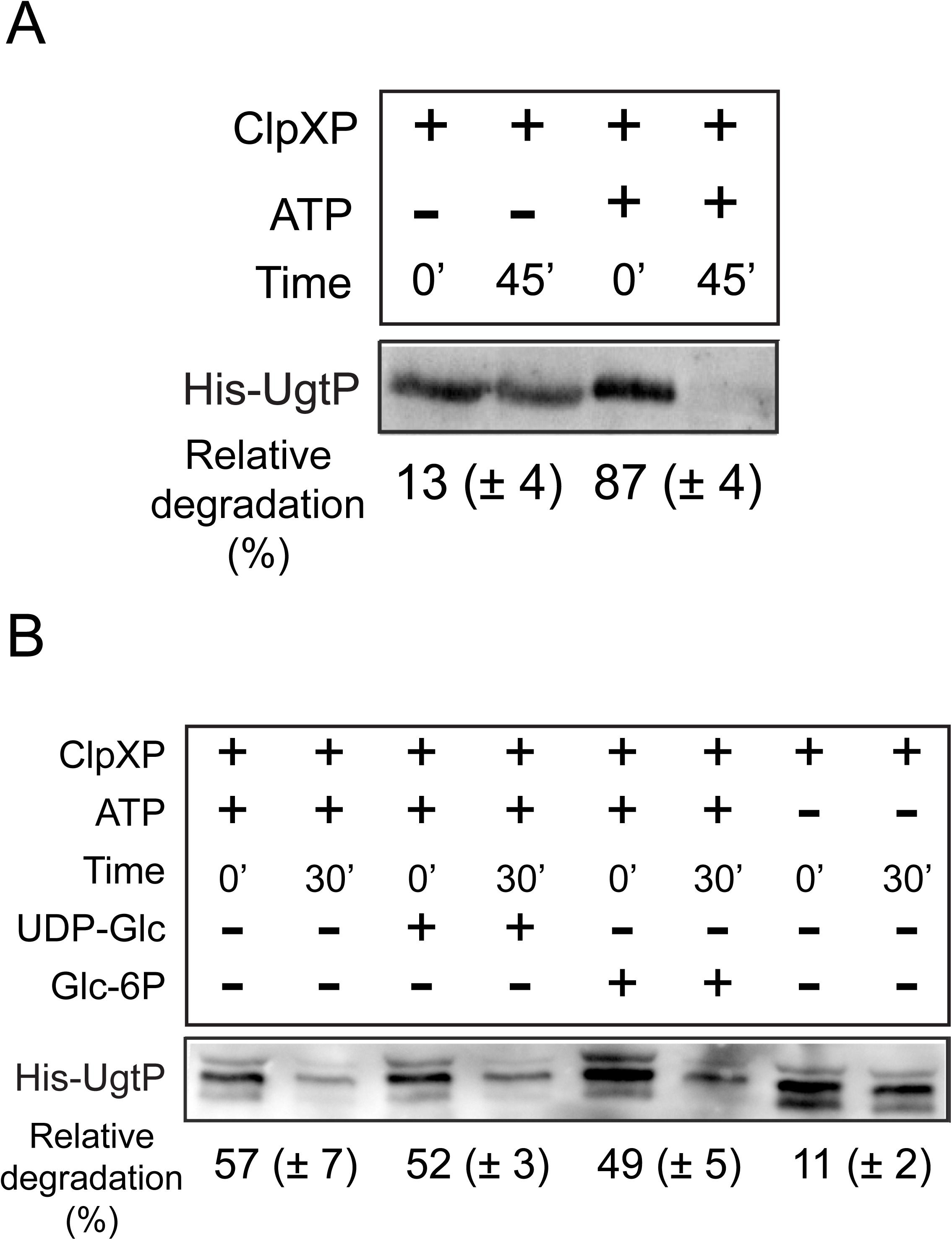
ClpXP targets UgtP for proteolysis *in vitro* independent of UDP-glucose. **(A)** Immunoblot of purified His-UgtP after incubation with purified ClpXP. Reactions consisting of 3μM ClpX, 6μM ClpP, 3μM His-UgtP, and 5mM ATP were incubated for 45 minutes at room temperature. ClpXP substrate controls are shown in (Additional File 6: Fig. S6). **(B)** In vitro ClpXP cleavage assay ± UDP-glucose and glucose-6P. The assay used 3μM ClpX, 6μM ClpP, 3μM His-UgtP, 5mM ATP, and 2mM of either UDP-glucose or glucose-6P. α-His was used to visualize His-UgtP levels by immunoblot. Each cognate set was used to gauge relative degradation (%) as shown below (n = 3, error = SD).

Based on our *in vivo* data supporting a model in which hexose binding may shield UgtP from Clp-mediated proteolysis (Fig. 4), we speculated that adding UDP-glucose or simply glucose to the ClpXP proteolysis assays might hinder UgtP degradation. To test this model we added UDP-glucose or glucose 6-phosphate in excess to the ClpXP *in vitro* proteolysis reactions described above. However, we saw no difference in UgtP proteolysis in the presence of either sugar (Fig. 5B).

### Clp-mediated UgtP degradation during growth in nutrient-poor medium does not significantly impact cell size or diglucosyl-diacylglycerol production

In an effort to illuminate the potential “rationale” for Clp mediated degradation of UgtP during growth in nutrient-poor medium we examined the impact of Clp activity on cell size and production of diglucosyl-diacylglycerol (DGD), the anchor for LTA. UgtP exhibits a high affinity for FtsZ (38nM) even in the absence of UDP-glucose (24). Nutrient-dependent degradation may thus serve as an additional control to prevent UgtP-mediated division inhibition during growth in nutrient-poor conditions.

To test this hypothesis, we took advantage of two strains capable of producing a range of UgtP concentrations in minimal sorbitol: BH736 (*ΔugtP; P*_*xyl*_*-ugtP-his)* and BH12 (*P*_*ugtP*_*-ugtP-his; P*_*xyl*_*-ugtP-his*). Quantitative immunoblots indicate that non-induced BH736 produces no detectible UgtP-His, induced BH736 and non-induced BH12 produce similar levels of UgtP-His, and induced BH12 produces approximately 60% more UgtP-His than BH12 non-induced (Additional File 7: Fig. S7). If cell size under nutrient-poor conditions is sensitive to UgtP concentration, then we would expect to see a gradient of cell sizes with cell size increasing in proportion to UgtP concentration. (We opted not to measure the impact of ClpP on cell size since *clpP* is highly pleotropic and deletion could feasibly alter cell length independent of UgtP degradation)

Somewhat surprisingly, we did not observe a significant difference in cell size between strains regardless of the presence of xylose. Average cell size was not significantly different between any set of conditions, including the UgtP-null (BH736 - xylose) and UgtP-overexpression conditions (BH12 + xylose) (Table 1, p = 0.28). Additionally, cell size distributions for each condition are not obviously different from one another (Fig. 6A). These data fail to support a model in which Clp-mediated UgtP degradation is important for maintaining cell size in nutrient-poor medium.

**Table 1.**
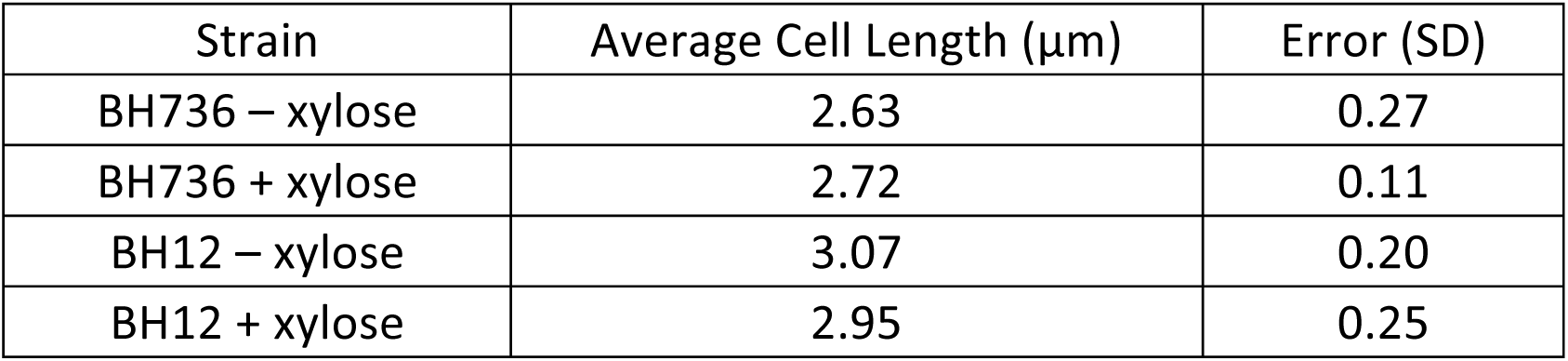
Average cell length in minimal sorbitol.

**Figure 6.**
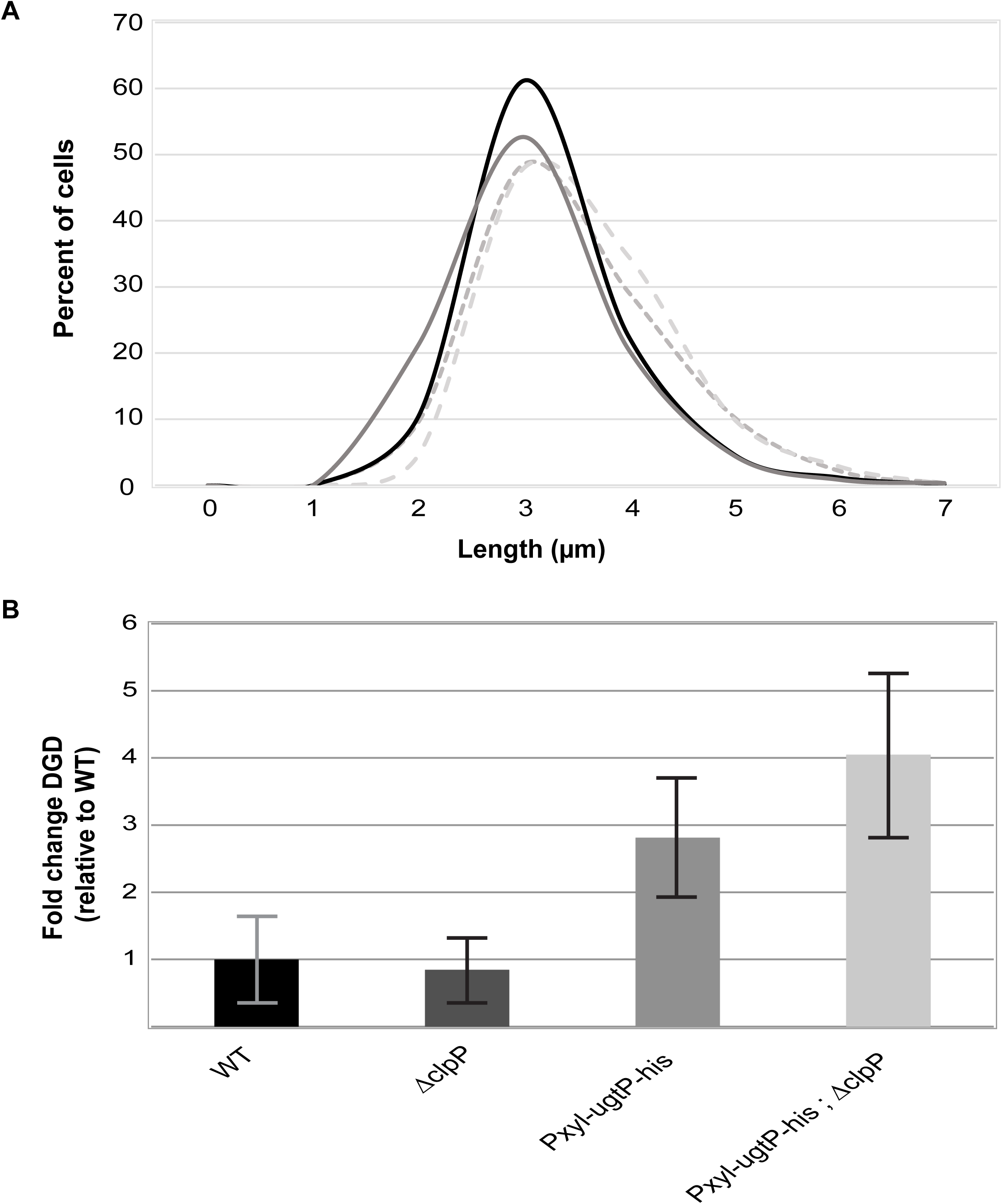
UgtP levels do not significantly affect cell length or diglucosyl-diacylglycerol levels in nutrient-poor media. **(A)** Cell length distributions of strains BH736 (*ΔugtP; P_xyl-ugtP-his_)* and BH12 (*P_*ugtP-ugtP-his*;_ P*_*xyl-ugtP-his*_) cultured in minimal sorbitol ± 0.5% xylose, determined by measuring the distance between the mid-points of adjacent cell wall septa (n = 600, error = SD). **(B)** Relative diglucosyl-diacylglycerol concentrations of lipid extracts from strains PL522 (WT), PL2102 (*ΔclpP*), BH10 (*P*_*xyl-ugtP-his*_), BH129 (*P*_*xyl-ugtP-his; ΔclpP*_) cultured in minimal sorbitol, determined by thin layer chromatography and subsequent densitometric analysis of separated lipids (n = 3, error = SD).

Average cell length of strains BH736 (*ΔugtP; P*_*xyl*_*-ugtP-his)* and BH12 (*P*_*ugtP*_*-ugtP-his; P*_*xyl*_*-ugtP-his*) cultured in minimal sorbitol ± 0.5% xylose, determined by measuring the distance between the mid-points of adjacent cell wall septa (n = 600, error = SD).

Another possible explanation for the degradation of UgtP in nutrient-poor conditions is the preservation of intracellular glucose by curtailing production of UgtP’s product, DGD (18). To test this possibility we measured DGD levels in wild type and Δ*clpP* strains, as well as their congenic *P*_*xyl*_*-ugtP* strains, cultured in minimal sorbitol (with xylose added when growing *P*_*xyl*_*-ugtP* strains). Briefly, we purified membranes from these strains, performed a methanol:chloroform lipid extraction, separated the lipids using thin-layer chromatography, stained for DGD using iodine gas, and quantified DGD using ImageJ. Lipids were also extracted from *ΔugtP* cells for use as a negative control.

Despite the 3-fold difference in UgtP concentration, DGD levels were not significantly different between WT/*ΔclpP* cells (p = 0.78) or induced *P*_*xyl*_*-ugtP* cells ± *clpP* (p = 0.30) (Fig. 6B). As expected, *ΔugtP* cells produced no detectible DGD (Additional File 8: Fig. S8). These data suggest that while UgtP is necessary for DGD production, degradation of UgtP via the Clp proteases is irrelevant for modulating DGD concentration. It is therefore unlikely that degradation of UgtP occurs in nutrient-poor conditions as a means of conserving glucose, as production of glucose-rich DGD is not affected by UgtP degradation.

## DISCUSSION

Our results indicate UgtP accumulation is controlled in a nutrient-dependent manner via a post-translational, Clp-dependent mechanism. This additional layer of control governing UgtP activity, distinct from UDP-glucose-mediated changes in UgtP’s affinity for itself and FtsZ, ensures that active UgtP accumulates only under nutrient-rich growth conditions when it antagonizes cell division. While there are several prominent examples of conditional proteolysis coupled to growth phase (13, 14, 33), there are only a handful of examples in which proteolysis has been linked to growth rate and/or nutrient availability. Two such examples are from the Narberhaus lab: 1) The DNA-binding replication inhibitor CspD is selectively proteolyzed by the Lon protease during growth in nutrient-rich medium (2, 34) and 2) LpxC, a deacetylase involved in lipid A biosynthesis, is degraded only when cultured at slower growth rates by the protease FtsH (35).

We were somewhat surprised by our finding that UgtP proteolysis *in vivo* is accomplished via the redundant activity of ClpCP, ClpEP, and ClpXP (Fig. 3B). While ZapC is recognized and degraded by ClpXP and ClpAP in *E. coli*, to our knowledge, UgtP is the first published example of a Clp substrate recognized by all the Clp chaperone proteins in *B. subtilis* (28). Further, we find that genes encoding all three *clp* chaperones and the clpP protease are expressed at higher levels during slower growth, which likely contributes toward the growth medium-dependent regulation of UgtP (Fig. 3D). It is noteworthy that ClpX, previously identified as a direct inhibitor of FtsZ assembly in *B. subtilis* independent of its role in proteolysis, also modulates the stability of an entirely different FtsZ inhibitor (36).

Since AAA+ proteases exhibit distinct substrate-binding repertoires with only minor overlap in target specificity (37), it is curious how and why all three Clp chaperones are able to target UgtP. These chaperones either recognize unstructured peptide sequence tags (known as degrons) within a client protein or adaptor proteins bound to target substrates. Although ClpXP is capable of degrading UgtP *in vitro* (Fig. 5), we were unable to identify putative degrons for any of the three Clp chaperone proteins within the UgtP primary sequence through comparison with other verified and putative B. subtilis Clp substrates (25, 38, 39). It thus remains an open question how ClpXP, ClpCP, and ClpEP each target UgtP for degradation.

UgtP degradation was enhanced *in vivo* by two point mutations (E306A N309A) in its putative hexose-binding site suggesting ligand binding might afford some protection from ClpP-specific proteolysis (Fig. 4). While there is precedent in both prokaryotic and eukaryotic systems of small molecule ligands governing proteolytic susceptibility (40-42), our data do not cohesively support that UDP-glucose shields UgtP from degradation. Neither defects in UDP-glucose biosynthesis (4) nor mutations disrupting the putative nucleotide binding site (ΔURA, F112A V117A) had an impact on UgtP accumulation (Fig. 4). These data suggest ligand binding is largely irrelevant to UgtP stability and the hexose-binding mutation (E306A N309A) may simply lead to a conformation that exposes a Clp recognition sequence. We are thus left with enrichment of the Clp proteases under nutrient-poor conditions as the most parsimonious explanation for UgtP degradation under these conditions (Fig. 3D).

## CONCLUSIONS

As a whole, our data point to UgtP being subject to an elaborate, and apparently over-engineered set of multilayered controls. Not only is it unclear why three Clp chaperones are required for UgtP degradation in minimal medium, it is also not readily apparent why a carbon-starved cell would spend energy to transcribe and translate *ugtP* only to immediately have it proteolyzed, consuming ATP at each step. We were unable to identify a phenotypic explanation for the nutrient-dependent degradation of UgtP. UgtP levels had no effect on cell size during growth in nutrient-poor medium and diglucosyl-diacylglycerol production was largely independent of ClpP (Fig. 6 and Table 1). Given the myriad of conditions under which *B. subtilis* is able to grow, it may very well be that we are looking in the wrong place and for the wrong phenotypes. Alternatively, UgtP may simply be one of a large cohort of proteins that have a shorter half-life during growth in nutrient-poor conditions (43). If so, growth medium-dependent changes in Clp protease activity may simply serve as a crude means of supporting *B. subtilis’* uncanny ability to rapidly adapt to a wide range of environments while at the same time, ensuring cells retain a relatively robust supply of biosynthetic building blocks even during growth in nutrient-poor conditions.

## MATERIALS AND METHODS

### Strains and general methods

All *B. subtilis* strains are derivatives of JH642 (44). Details of their construction and a list of strains used for this study are described in (Additional File 9: Supplemental Materials and Methods and Additional File 10: **Table S1**). All cloning was done using the *E. coli* strain AG1111 (45). Either Vent (NEB) or Phusion (NEB) DNA polymerases were used for PCRs. Cells were cultured in either Lysogeny Broth (LB) or minimal S7_50_ defined media (46) supplemented with either 1% glucose, glycerol, or sorbitol as carbon sources and appropriate amino acid supplements. For strains containing *ΔclpP* or *ΔclpCEX*, cells were always first cultured in LB prior to dilution.

### Quantitative immunoblotting

Bacterial cells cultured to A_600_ 0.20-0.40 were lysed by centrifuging 1 mL of culture then re-suspending in 50 μL lysis buffer (20 mM tris pH 8.0, 12.5 mM MgCl_2_, 1 mM CaCl_2_, 2 mg/mL lysozyme, 1X Halt™ Protease Inhibitor Cocktail), incubating 10 minutes at 42°C, then adding SDS to 1% (v/v). Laemmli buffer was added to 1X, lysates were incubated 10 minutes at 100°C, then 5 μL lysate was subjected to SDS-PAGE using precast gels (Bio-Rad # 4561045). Proteins were transferred to PVDF membranes using Bio-Rad Trans-Blot^^®^^ Turbo™ instrument as described on pg. 15 of the operation manual. Total protein was stained using 1X Ponceau S solution in 5% acetic acid for 5 minutes. Membranes were then blocked with 5% nonfat milk in PBS for 1 hr.

Mouse monoclonal THE™ His Tag (Genscript) α-His antibodies incubated (at 1:1000) were used to detect His-tagged UgtP. Rabbit polyclonal (Genscript) α-GFP antibodies incubated (at 1:1000) were used to detect YFP-tagged UgtP. FtsZ was detected using an affinity-purified polyclonal rabbit α-FtsZ antibody (at 1:5000) (47). Cognate goat α-rabbit or goat α-mouse (Genscript) HRP secondary antibodies were used (at 1:5000). The membranes were developed using ECL substrate (Bio-Rad #1705060) and imaged using a Li-COR Odyssey FC instrument. Blot density was quantified using ImageJ (NIH) and processed in Microsoft Excel.

### *In vivo* UgtP degradation assay

Protocol was adapted from (29). Strains were cultured in 5 mL LB + 0.5% xylose at 37 °C with shaking until A_600_ = 0.20-0.40. These cultures were then diluted into 20 mL minimal sorbitol + 0.5% xylose to A_600_ = 0.005. Once A_600_ reached 0.20-0.40, spectinomycin was added to 200 μg/mL. Cultures were sampled (1 mL) at 0, 30, 60, 90, and 120 minutes after spectinomycin addition, and the samples were frozen at −80°C for future use. These samples were then subjected to quantitative immunoblotting for UgtP-His, the protein was quantified using ImageJ (NIH) and processed in Microsoft Excel.

### β-galactosidase activity in liquid cultures

Specific activity was calculated essentially as described in (48). Strains encoding *lacZ* fusions were cultured at 37°C to early/mid-log (A_600_ 0.30-0.40). Prior to sampling, the cultures were diluted 1:2 in their respective medium and absorbance at A_600_ was recorded. 30μl of toluene and 30μl of a 0.1% sodium dodecyl sulfate solution were added to 2ml of bacterial culture to permeabilize cells. Incubating cells at 37°C for 45 minutes then evaporated the toluene. Cells were then mixed with Z-buffer (60 mM Na_2_HPO_4_, 40 mM NaH_2_PO_4_, 10 mM KCl, 1 mM MgSO_4_, 50 mM β-mercaptoethanol) and tubes were incubated at 25°C for 5’. Reactions were started by adding 0.25 ml of 0.4% o-nitrophenol-β-galactoside in Z-buffer and stopped by adding 0.5 ml of 1 M Na_2_CO_3_. A_420_ was then recorded. The specific unit value was calculated using the equation: = 200 × (A_420_ of the culture - A_420_ in the control tube)/minutes of incubation × dilution factor.

### Quantitative real-time polymerase chain reaction

RNA was harvested from *B. subtilis* cells in early-log (A_600_ = 0.20-0.40) with RiboPure^™^ Kit (Ambion), treated with the Turbo DNA-Free Kit™ (Ambion), and reverse transcribed for 1 hour at 42°C using the RETROscript^®^ Kit (Ambion). Template was diluted 10-fold, added to iTaq SYBR Green Supermix (Bio-Rad) with appropriate oligonucleotide pairs. Oligonucleotides used in this study are described in (Additional File 11: **Table S2**). Data were acquired using an Applied Biosystems model 7500 thermocycler. Results were analyzed using the comparative Pfaffl method (49).

### *In vitro* ClpXP proteolysis assay

Reactions were carried out in 50mM HEPES pH 7.6, 50mM KCl, 15mM Mg acetate, 5mM DTT, 5mM ATP, 10mM creatine phosphate, and 0.1μg/μl creatine kinase (Sigma). 500ng of His-UgtP was incubated at 37°C in the presence of ClpP (6μM) and ClpX (3μM) in a 100μl reaction mixture. Each reaction was initiated by the addition of 5mM ATP. At 0 and 30 or 45 minutes, samples of the reactions were taken containing what corresponded to 125ng His-UgtP at the start of the reaction. Samples were then diluted to 2.5ng/μl with 2X sample buffer. 50ng of His-UgtP was then loaded per lane on a 10% SDS-PAGE. After electrophoresis, proteins were transferred via Western blot to a PVDF membrane. Membranes were blocked, then incubated with mouse monoclonal THE™ His Tag (Genscript) at a 1:4000 dilution overnight at 4°C. Control reactions with Spx and Thio-His were performed as with His-UgtP except at a concentration of 3μM. At 0 and 60 minutes, 15μl of the samples was collected and analyzed on a 15% SDS-PAGE followed by staining with Coomassie blue.

### Fluorescence microscopy and cell length measurements

Samples were first fixed as previously described (50). Briefly, cells were fixed by treating with 2.575% paraformaldehyde and 0.0008% gluteraldehyde. Fixed samples were permeabilized with 0.01 mg/mL lysozyme for two minutes, and then treated with 10 μg/mL wheat germ agglutinin tetramethylrhodamine conjugate (WGA-Rhod, ThermoFisher #W849) in PBS to stain cell wall septa for 10 minutes. 5 μL of prepared samples were applied to 1% agarose in PBS pads, allowed to dry, and then covered with a coverslip. Cells were imaged using a Nikon TI-E microscope equipped with a Texas Red/mCherry/AlexaFluor 594 filter set for fluorescence microscopy (Chroma # 39010). Cell length was determined using Nikon Elements by measuring the distance between the midpoints of adjacent cell wall septa for cells stained with WGA-Rhod. T-test analysis was employed to test for significant differences between conditions.

### Lipid Extraction

Bacterial cultures (1L volumes) were centrifuged at 15,000 x g for 1 minute, decanted, and then resuspended in 100 mL 100 mM sodium citrate pH 4.7. Cells were lysed using a French press. Lysates were centrifuged at 12,000 g for 20 minutes to pellet membranes. Pellets were weighed and resuspended to 0.4 g/mL in 100 mM sodium citrate pH 4.7. Methanol and chloroform were added to obtain a ratio of 2:1:0.8 methanol:chloroform:sodium citrate. Mixture was vortexed 30 seconds every 15 minutes for 2 hours. Chloroform and sodium citrate were added to obtain a ratio of 1:1:0.9 methanol:chloroform:sodium citrate. The mixture was vortexed for 1 minute and centrifuged at 4000 x g for 10 minutes. Bottom chloroform layer was transferred to a new tube. Methanol and sodium citrate were added to obtain a ratio of 1:1:0.9 methanol:chloroform:sodium citrate. Mixture was vortexed for 1 minute and centrifuged at 4000 x g for 10 minutes. Bottom chloroform layer was transferred to new tube and allowed to dry in fume hood. Dried lipids were weighed and resuspended in 1:1 methanol:chloroform to a final concentration of 600 mg/mL and stored at −20 °C.

### Lipid Thin-Layer Chromatography and diglucosyl-diacylglycerol quantification

Added 140 mL chloroform and 60 mL methanol to a glass TLC development chamber. Placed filter paper into developing solution, thoroughly wetted blot paper, and then placed the paper along the chamber walls. A TLC plate (Analtech #p46021) was pre-run in developing solution by placing it in the chamber, covering the chamber with its glass lid, and allowing the solvent to run to the top of the plate. The plate was dried at 100 °C for 10 minutes. Using a pencil, a line was drawn across the width of the plate approximately 1 cm from the bottom. Lipid samples (2μL) were spotted along this line and allowed to dry. Samples were repeatedly spotted and allowed to dry 5 additional times. Once the spots were dry, the plate was placed back into the chamber, the lid was placed on top, and the solvent was allowed to run to 1cm from the top of the plate. The plate was taken out of the development chamber, the solvent line was quickly marked with pencil, and the plate was dried. The plate was then placed inside a polystyrene container with a few iodine crystals, sealed, and stained for 16 hours with iodine gas. The stained plate was scanned to acquire images. The stained plate was scanned to acquire images. ImageJ was used to quantify the amount of DGD present, and another band (indicated in Additional File 8: Fig. S8) was used as a loading control. T-test analysis was employed to test for significant differences between conditions.

## LIST OF ABBREVIATIONS

LB: Lysogeny Broth
LTA: Lipoteichoic Acid
OPG: Osmoregulated Periplasmic Glucan
Minimal Glucose: S7_50_ + 1% Glucose
Minimal Glycerol: S7_50_ + 1% Glycerol
Minimal Sorbitol: S7_50_ + 1% Sorbitol
DGD: diglucosyl-diacylglycerol

## ADDITIONAL FILES

**Additional File 1** – Adobe Acrobat .pdf format, YFP-UgtP is degraded in nutrient-poor growth conditions, this file shows immunoblots of ectopically expressed YFP-UgtP cultured in nutrient-rich and nutrient-poor media.

**Additional File 2 –** Adobe Acrobat .pdf format, representative immunoblot for *in vivo* UgtP degradation experiment, this file shows an immunoblot of UgtP-His from cells with and without *clpP*, cultured in nutrient-poor media over a 2-hour period, after addition of spectinomycin.

**Additional File 3 –** Adobe Acrobat .pdf format, the UgtP oligomer mutant has WT localization and delays cell division in a *ΔpgcA* background, this file shows immunofluorescent microscopic images of cells harboring either *yfp-ugtP* or *yfp-ugtP(ΔOLI)* in a *ΔpgcA* background cultured in nutrient-rich media, and also shows cell length distributions for the previously mentioned strains.

**Additional File 4 –** Adobe Acrobat .pdf format, *ugtP(ΔHEX)* is expressed at the same level as *ugtP* and UgtP(ΔHEX) is stabilized in a *ΔclpP* background, this file shows qRT-PCR data for UgtP binding mutants compared to WT UgtP, and also shows quantitative immunoblots for UgtP-His from the previously mentioned strains cultured in both nutrient-rich and nutrient-poor media.

**Additional File 5 –** Adobe Acrobat .pdf format, a comparison of UDP-glucose molecules per cell during nutrient-rich and -poor conditions, this file shows UDP-glucose molecules per cell from WT, *ΔpgcA*, and *ΔugtP* cells cultured in nutrient-rich media (and WT in nutrient-poor media) as measured by LC-MS/MS.

**Additional File 6 –** Adobe Acrobat .pdf format, positive and negative controls for the ClpXP *in vitro* proteolysis assay, this file shows proteolysis of both a known substrate for ClpXP (Spx), and a non-targeted protein (Thioredoxin-His).

**Additional File 7 –** Adobe Acrobat .pdf format, UgtP concentration can be modulated in nutrient-poor media, this file shows quantitative immunoblots for UgtP-His from strains containing either one inducible copy of *ugtP-his*, or one inducible copy and one “native” copy of *ugtP-his*, cultured in nutrient-poor media.

**Additional File 8 –** Adobe Acrobat .pdf format, representative TLC of membrane lipids, including DGD, this file shows lipid extracts from strains producing variable amounts of UgtP cultured in nutrient-poor media, separated on a TLC plate.

**Additional File 9 –** Adobe Acrobat .pdf format, supplemental materials and methods, this file contains materials and methods for the experiments performed in additional files 1-8.

**Additional File 10 –** Adobe Acrobat .pdf format, bacterial strains used in this study, this file contains a table of strains, their genotypes, and their sources used in this study.

**Additional File 11 –** Adobe Acrobat .pdf format, oligonucleotide sequences used for qRT-PCR, this file contains a table of the oligonucleotide sequences used for qRT-PCR of various genes.

**Additional File 12 –** Adobe Acrobat .pdf format, supplemental references, this file contains references for the materials/methods and tables from **Additional Files 9-11**

## DECLARATIONS

### Ethics Approval and Consent to Participate

Ethics approval was not required for this study, which utilized only bacteria and did not involve humans, human data, or animals.

### Consent for Publication

Consent for publication was not required for this study, which utilized only bacteria and did not involve humans, human data, or animals.

### Availability of Data and Materials

The datasets used and/or analyzed during the current study are available from the corresponding author on reasonable request.

### Authors Contributions

NH designed and performed all aspects of the experimentation except as noted below, analyzed data, and drafted the manuscript. JZ performed and analyzed data for the *in vivo* UgtP-His proteolysis assay in (Fig. 3C), cell measurements and membrane lipid TLC in (Fig. 6 and Table 1), Western blots of YFP-UgtP in (Additional File 1: Fig. S1), Western blots to demonstrate differences in UgtP-His expression in (Additional File 7: Fig. S7). JZ also designed the cell measurement experiment in (Fig. 6A and Table 1) and assisted in drafting the manuscript. PB performed, designed, and analyzed the *in vitro* His-UgtP proteolysis assay in (Fig. 5 and Additional File 6: Fig. S6) including the purification of ClpXP and UgtP. PB also assisted in drafting the manuscript. AC constructed the hexose/uracil/oligomerization mutant forms of *ugtP* used in (Fig. 4 and Additional File 4: Fig. S4). PAL helped design experiments and draft the manuscript. All authors read and approved the final manuscript.

## ACKNOWLEDGEMENTS

We are indebted to Richard Losick, Peter Zuber, and David Dubnau for the kind gift of strains and reagents. Finally, we thank all members of the Levin lab for thoughtful discussions. This work was supported by a National Institutes of Health grant (NIH GM64671) to PAL.

---

**Additional File 1:**
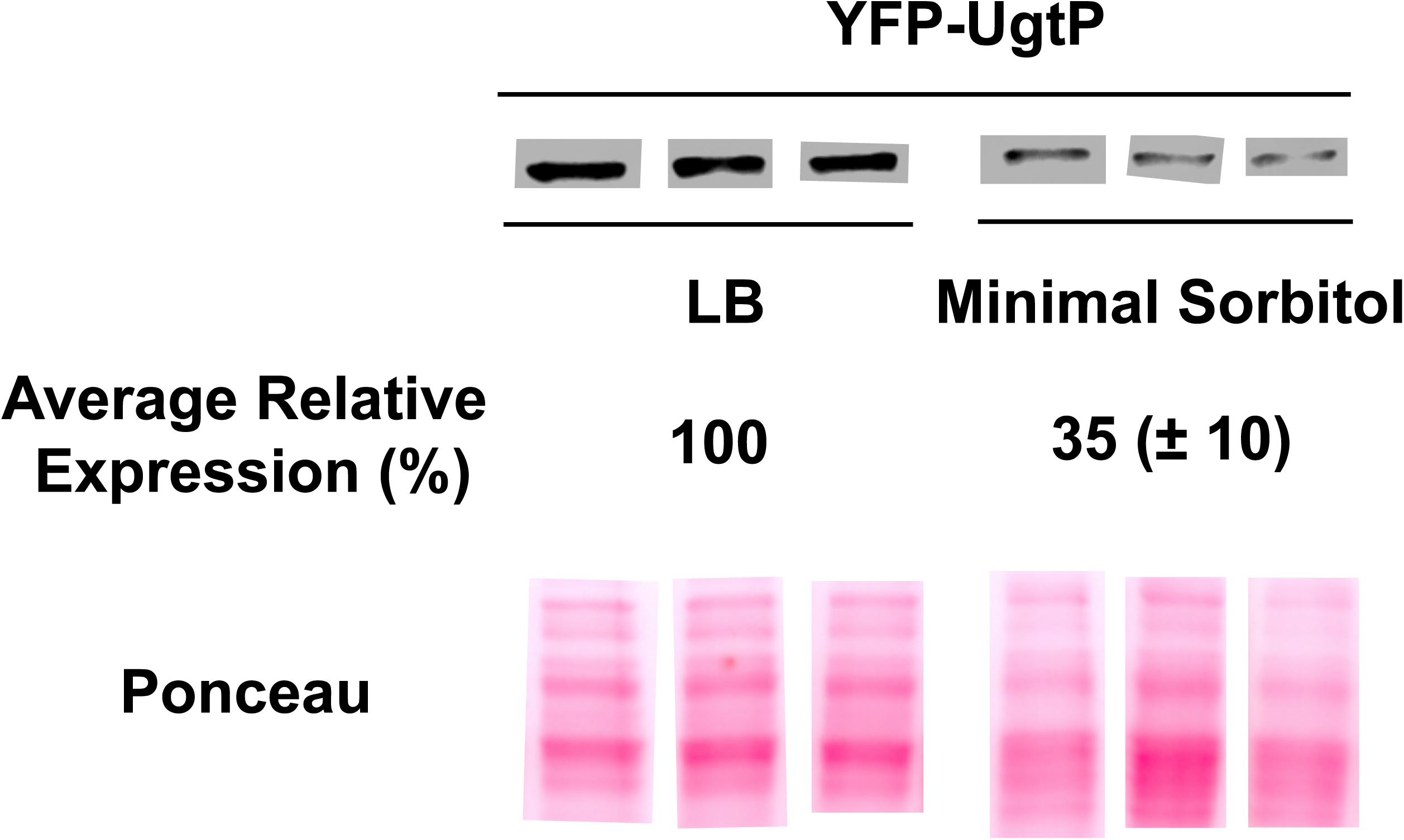
YFP-UgtP is degraded in nutrient-poor growth conditions. Quantitative immunoblot of ectopically expressed YFP-UgtP in nutri-ent-rich and nutrient-poor growth conditions. PL2423 (*Pxyl-yfp-ugtP*) was grown in both LB and minimal sorbitol medium. Total protein was used as a loading control via Ponceau staining. Protein levels in LB are set as the reference in the relative expression values shown (n=3, error = SD).

**Additional File 2:**
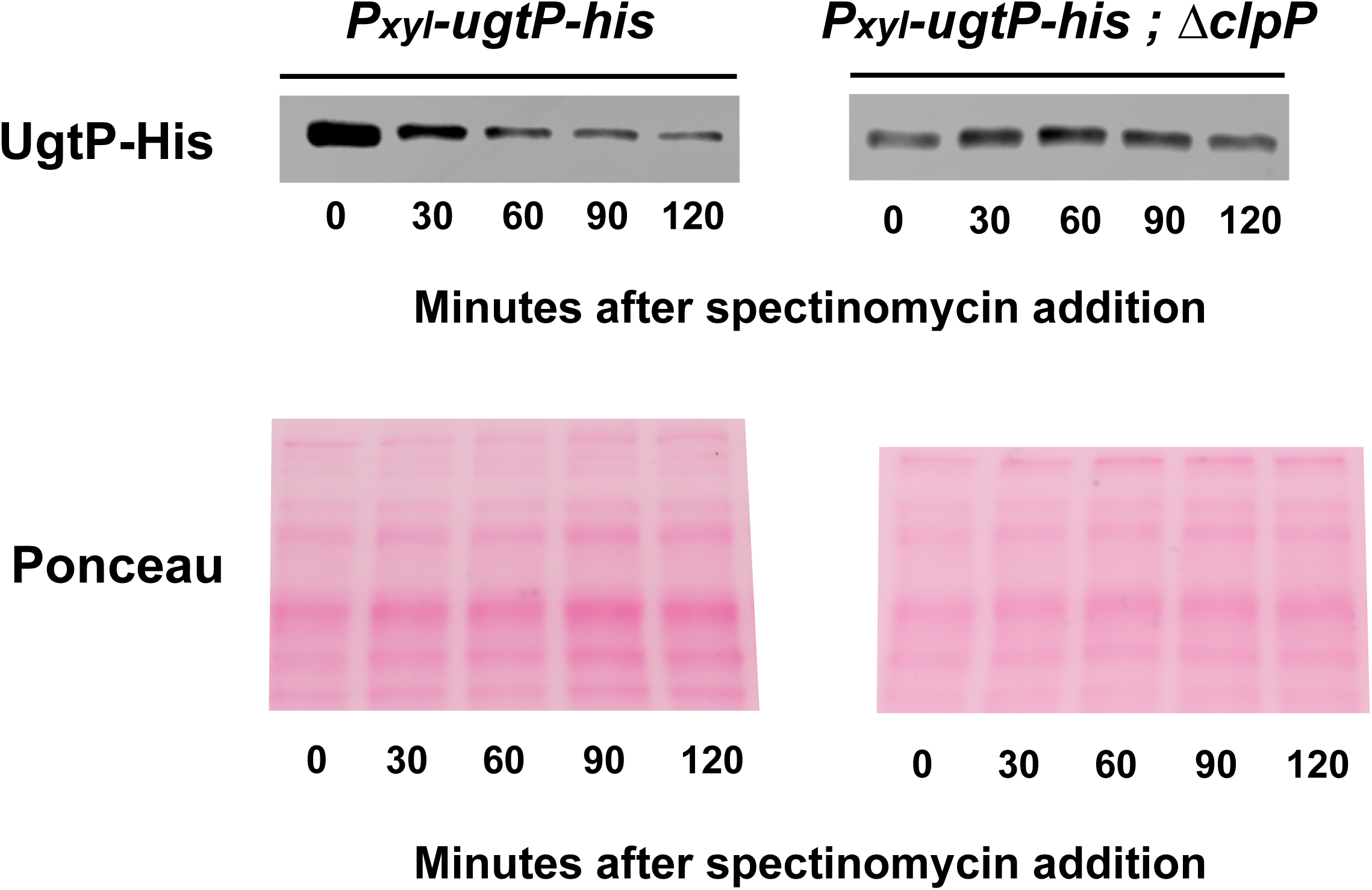
Representative immunoblot for *in vivo* UgtP degradation experiment. Representative immunoblot of UgtP-His from both BH10 (*Pxyl-ugtP-his*) and BH129 (*Pxyl-ugtP-his; ΔclpP*) cultured in S750 sorbitol + 0.5% xylose. Spectinomycin was added to cells to inhibit new protein synthesis, and then cultures were sampled every 30 minutes for 2 hours. Ponceau staining was performed as a loading control.

**Additional File 3:**
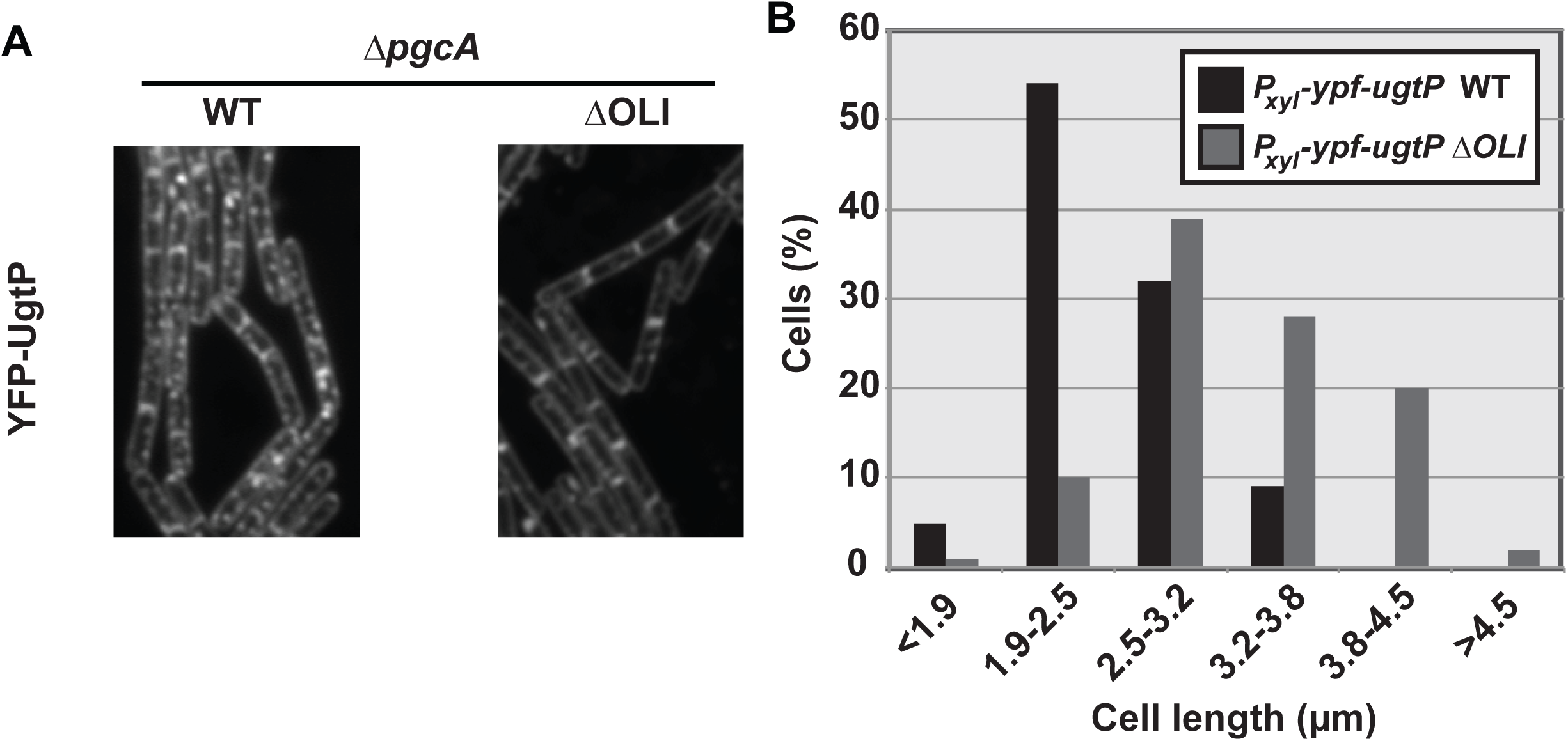
The UgtP oligomer mutant has WT localization and delays cell division in a *ΔpgcA* background. **(A)** WT YFP-UgtP (PL2292: *Pxyl-yfp-ugtP*) in cells unable to synthesize its substrate UDP-glucose (*ΔpgcA* background) self associates into punctate foci in LB + 0.5% xylose (1). However, mutating the putative oligomerization site (JC395: *Pxyl-yfp-ugtP* I142A E146A) substantially reduces the number of UgtP puncta to localize with the septum. Scale bar = 3 μm. **(B)** A histogram of *Pxyl-yfp-ugtP* (PL2292) or *Pxyl-yfp-ugtP ΔOLI* (JC395) cell length when grown in LB + 0.5% xylose (n > 150 measurements per strain).

**Additional File 4:**
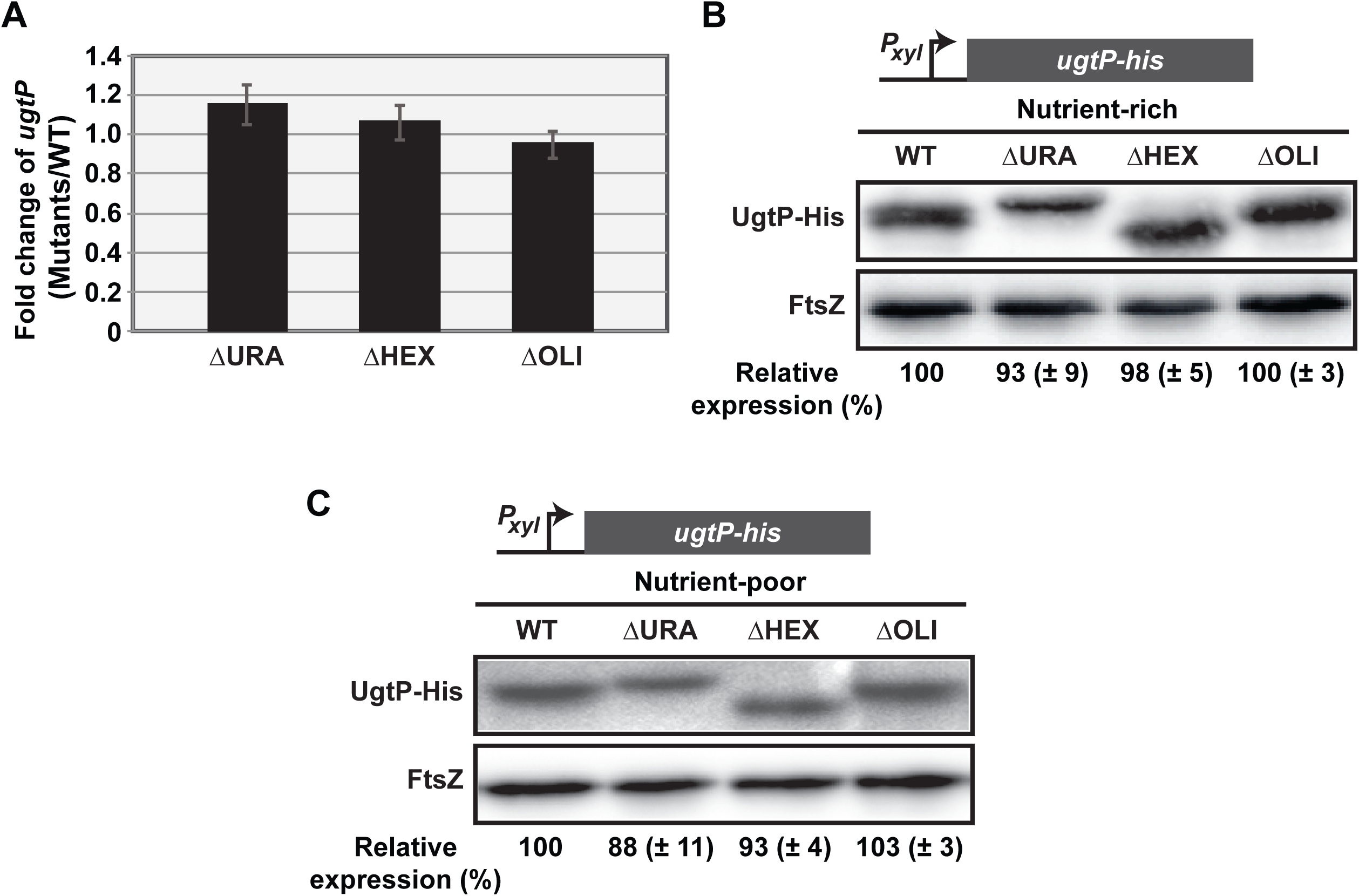
*ugtPΔHEX* is expressed at the same level as *ugtP* and UgtPΔHEX is stabilized in a *ΔclpP* background. **(A)** *ugtP* transcript levels of WT versus the ΔURA, ΔHEX, and ΔOLI mutants cultured in LB + 0.5% xylose measured by qRT-PCR (n = 3, error = SD). The *Pxyl-ugtP-his* variants in a *ΔclpP* background (BH767, BH769, BH771, BH773) grown in either **(B)** LB + 0.5% xylose or **(C)** minimal sorbitol + 0.5% xylose. FtsZ is used as a loading control. The relative expression (%), using WT UgtP as the reference, is shown below (n = 3, error is SD).

**Additional File 5:**
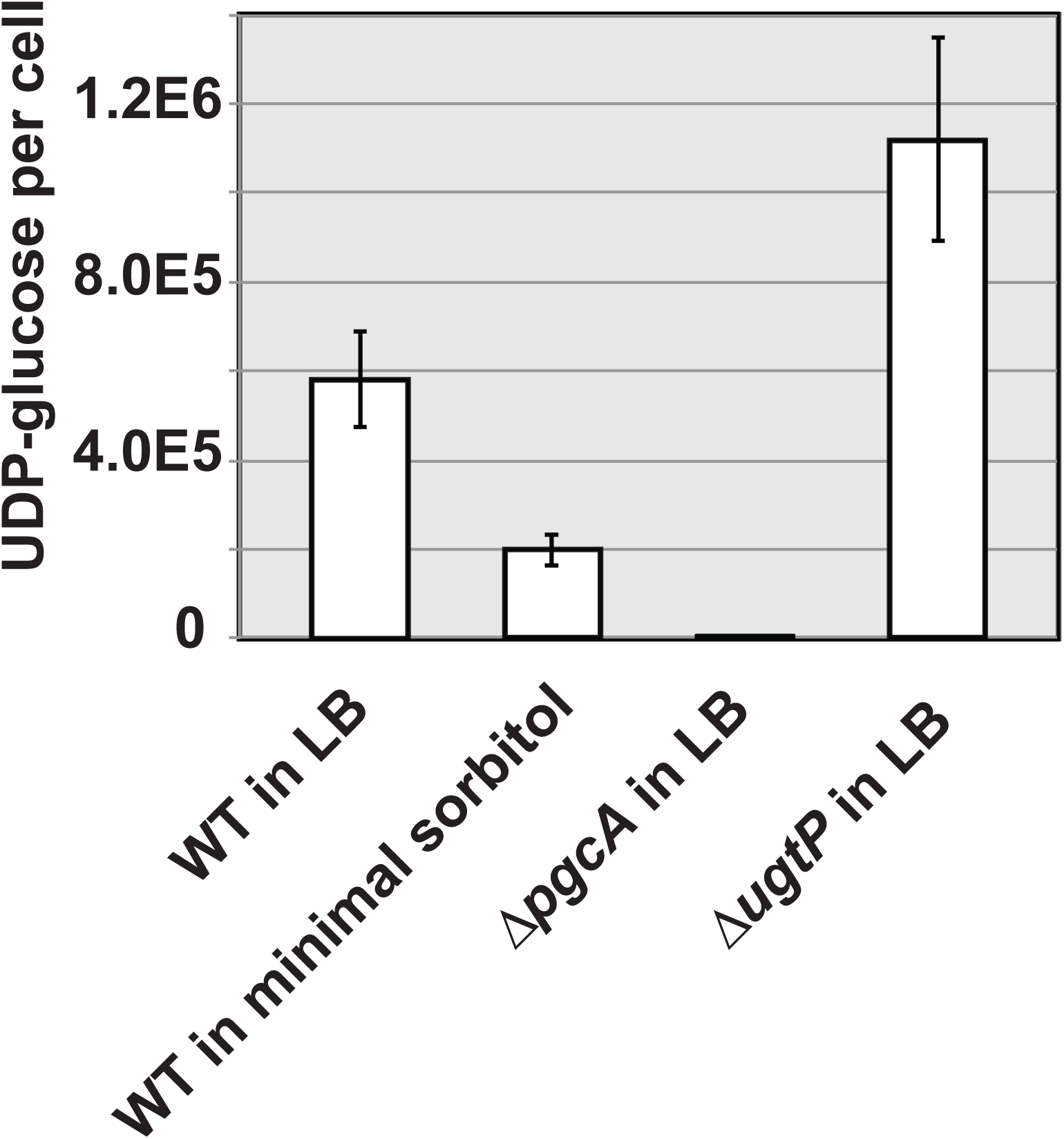
A comparison of UDP-glucose molecules per cell during nutrient-rich and -poor conditions. The intracellular UDP-glc levels of WT B. subtilis (PL522) grown in either LB or minimal sorbitol were measured by LC-MS/MS. Appropriately, when cultured in LB we observed reduced UDP-glucose levels in the *ΔpgcA* mutant (PL1310) (a phosphoglucomutase upstream in the UDP-glucose metabolic pathway), while knocking out *ugtP* (PL2265) increased UDP-glucose. (n = 3, error bars = SD.)

**Additional File 6:**
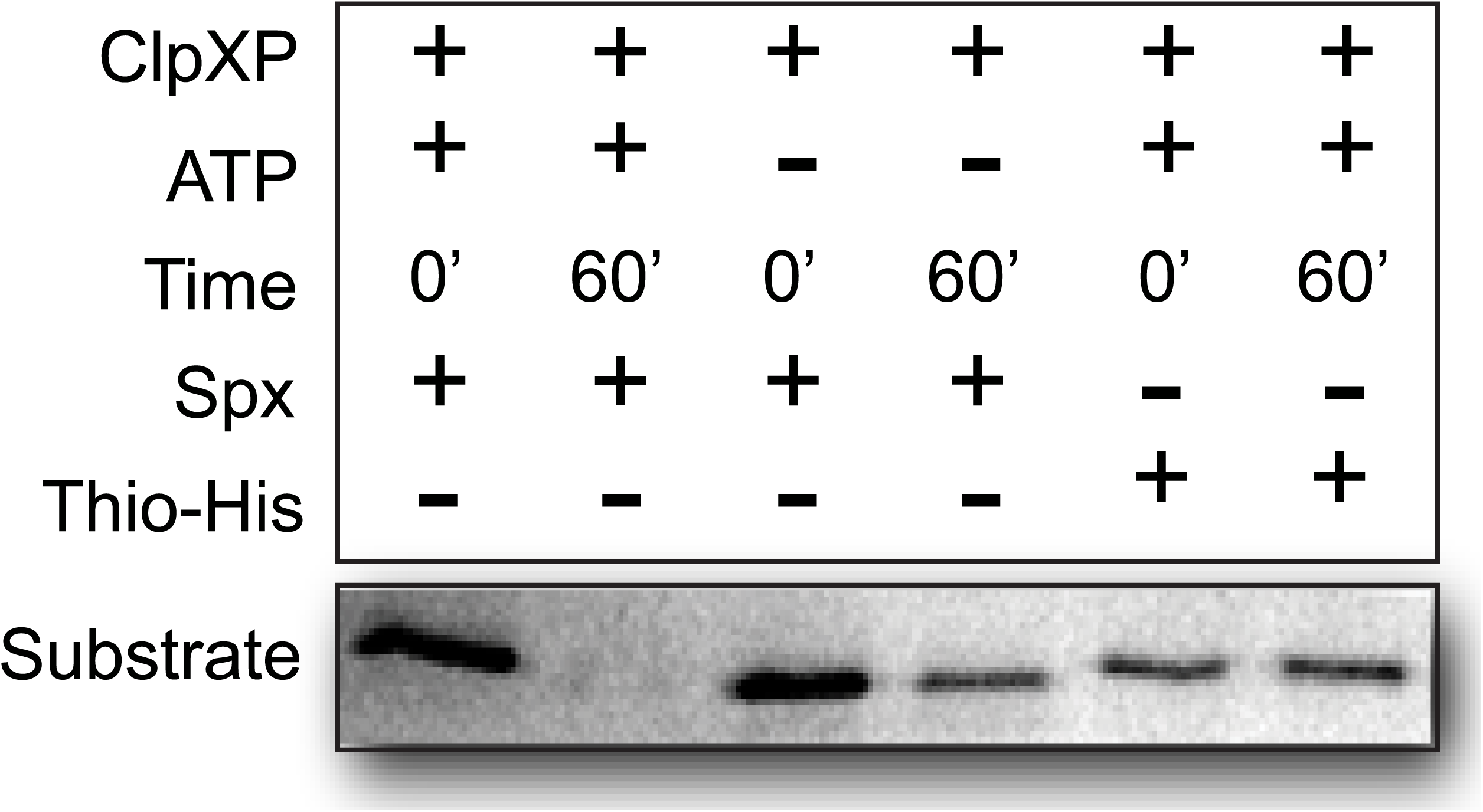
Positive and negative controls for the ClpXP *in vitro* proteolysis assay. A Coomassie stained gel examining ClpXP-mediated proteolysis of a known substrate, Spx (15.5 kDa), and a non-targeted protein, Thioredoxin-His (17.1 kDa). A mixture of 3μM ClpX, 6μM ClpP, 5mM ATP, and either 3μM of Spx or Thio-His was incubated for 60 minutes.

**Additional File 7:**
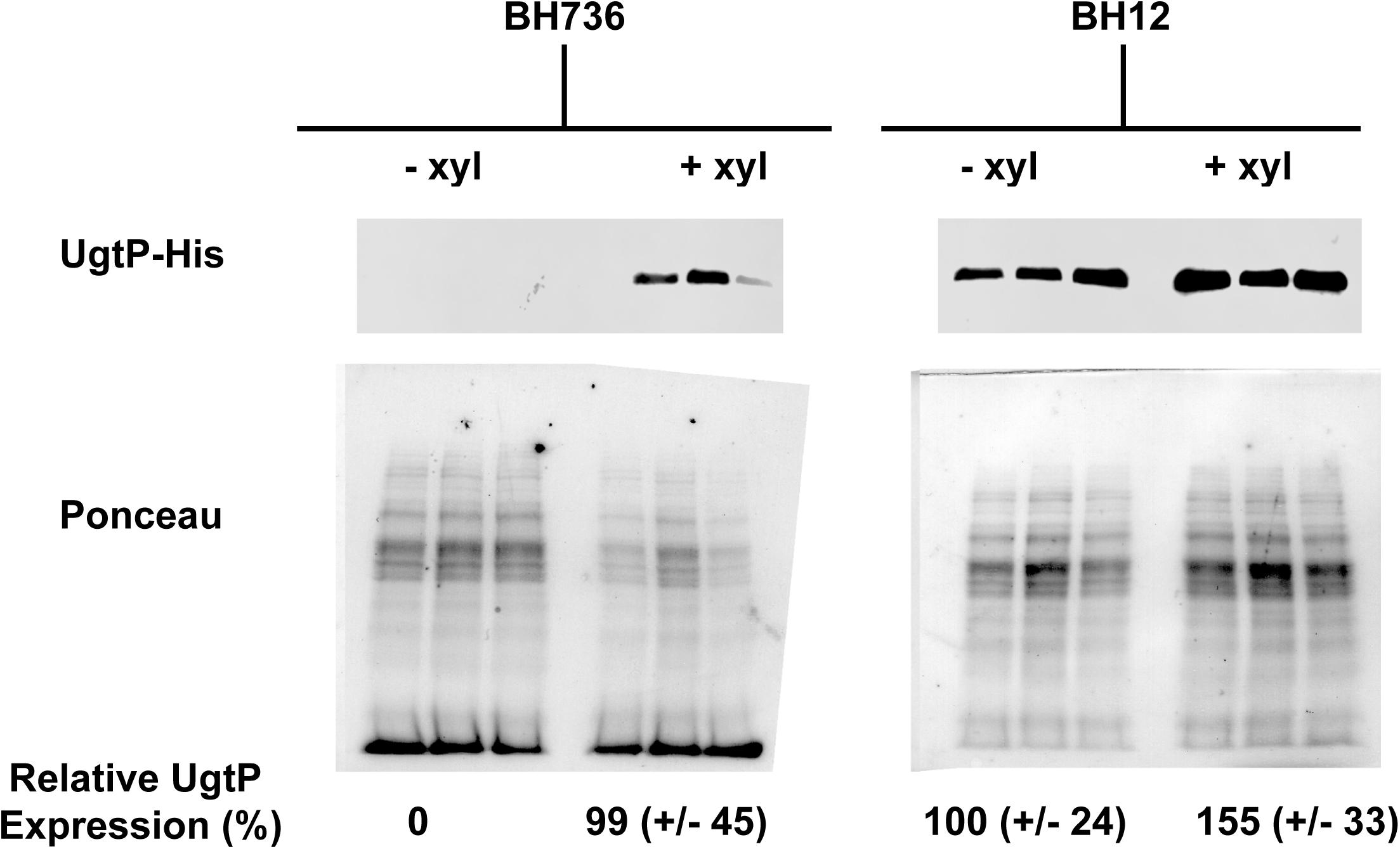
UgtP concentration can be modulated in nutri-ent-poor media. Western blots of UgtP-His from BH736 (*ΔugtP; Pxyl-ugtP-his*) and BH12 (*PugtP-ugtP-his; Pxyl-ugtP-his*) cells cultured in minimal sorbitol, with and without xylose, using total protein as a loading control via Ponceau staining. Protein levels of BH12 - xylose are set as the reference in the relative expression values shown, as they represent WT expression (n = 3, error = SD).

**Additional File 8:**
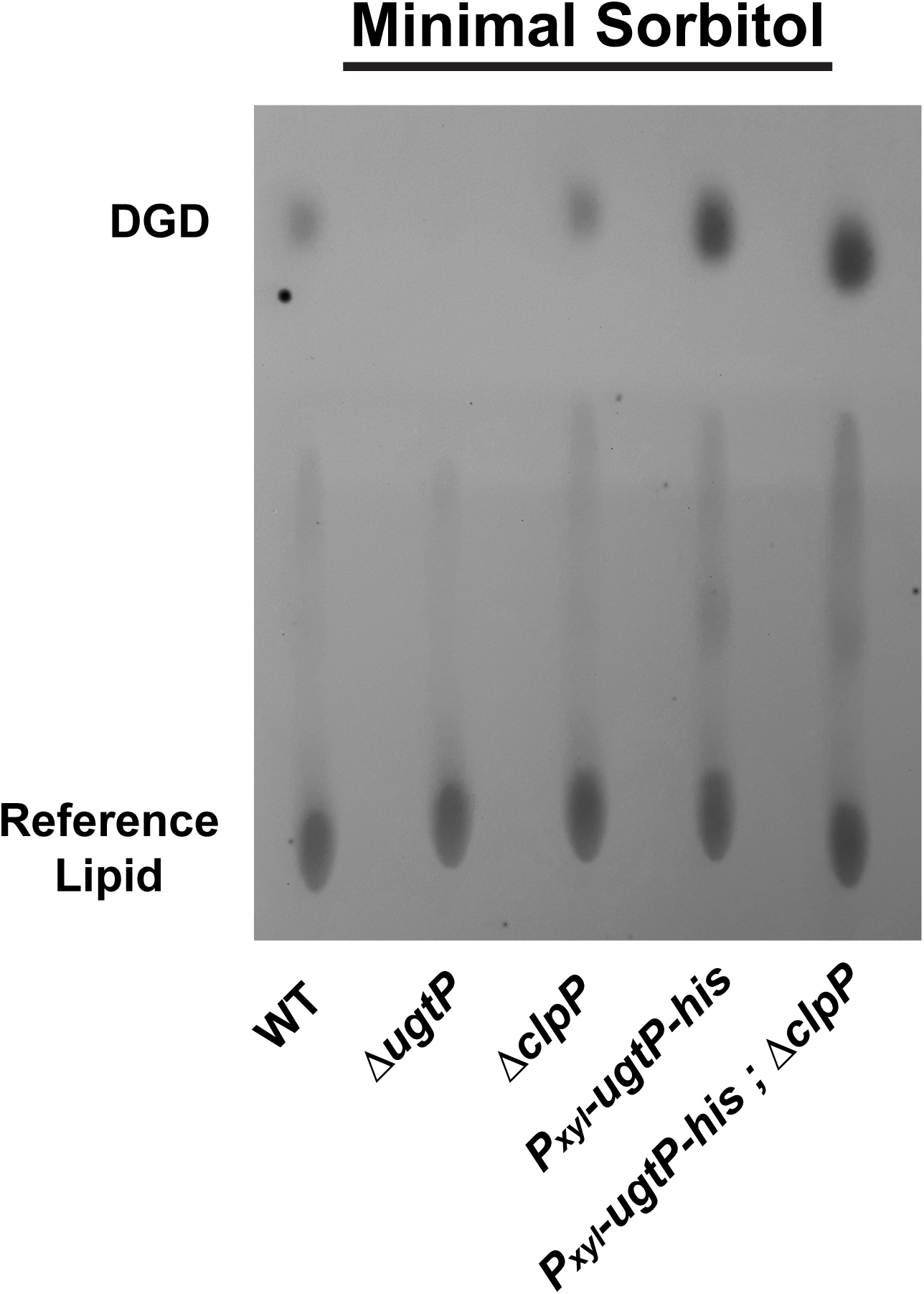
Representative TLC of membrane lipids, including DGD. Strains PL522 (WT), PL2268 (*ΔugtP*), PL2102 (*ΔclpP*), BH10 (*Pxyl-ugtP-his*) and BH129 (*Pxyl-ugtP-his; ΔclpP*) were grown in minimal sorbitol prior to membrane lipid isolation via methanol/chloroform extraction and TLC. TLC plates were developed by staining with gaseous

## Additional File 9: Supplemental Materials and Methods

### Strain construction

Ampicillin was used at 100μg/ml, chloramphenicol at 5μg/ml for *B. subtilis* and 30μg/ml for *E. coli*, kanamycin at 5μg/ml for *B. subtilis* and 50μg/ml for *E. coli*, tetracycline at 12.5μg/ml, and spectinomycin at 100μg/ml. MLS resistance was selected for using erythromycin at 0.5μg/ml and lincomycin at 12.5μg/ml. Tryptophan and phenylalanine were supplemented at 40μg/ml, threonine at 80μg/ml.

The *PugtP-lacZ* transcriptional fusion was built by amplifying ∼700bp of the *ugtP* promoter region using oligos: AGTCGAATTCGCAGGTTTGTATTACCATTACG and AGTCGGATCCAATGTAATCAACAACAAG. The translational fusion additionally encoded the first 90 bases of *ugtP* and was amplified with oligos: AGTCGAATTCGCAGGTTTGTATTACCATTACG and AGTCGGATCCATGTGTAACAAGTATTTCACAAAACCG. The PCR products and the vector pDG268 were digested with BamHI and SphI and ligated together. This construct was then transformed into PL522 facilitating the integration of *PugtP-lacZ* at the amyE locus.

The *Pxyl-ugtP-his* was built by amplifying *ugtP-his* from PL2265 chromosomal DNA using oligos: ATAGCATGCACATTGAGGTGAATTTACTTGAATACC and AGTCGGATCCCGATAGCACTTTGGCTTTTTG. The resulting PCR product and vector pRCD19 were digested with SphI and BamHI, then ligated together. This construct was then transformed into *B. subtilis*, which resulted in the integration of *Pxyl-ugtP-his* into the *thrC* locus.

The *his-ugtP* used in the ClpXP proteolysis assay was PCR amplified from PL522 genomic DNA using the following primers: GAGATCCATATGAATACCAATAAAAGAGTATTAATTTTGACTG and GATCGGATCCTTACGATAGCACTTTGGCTTTTTGTTTG. The *ugtP* gene product and pET28a(+) were digested with NdeI and BamHI and ligated. The resulting plasmid yielded a 6X-His tag with the linker sequence SSGLVPRGSH fused to the N-terminus of UgtP (PL3521).

The uracil, hexose, and oligomerization ugtP mutants were made using site-directed mutagenesis. The *Pxyl-ugtP-his*/pRCD19 plasmid was amplified using Phusion HF DNA polymerase (NEB) with the following oligos and its complement (not shown) (mutations underlined): uracil-binding mutant CCCGATATTATTATTAATACA<u>GCC</u>CCGATGATC<u>GCC</u>GTGCCGGAATACAG, hexose-binding mutant CCCGTGCCTGGACAGG<u>T</u>AAAAGAA<u>GCA</u>GCAAACTTCTTTGAAG oligomerization mutant GTCTTCATAAA<u>GC</u>TTGGGTTCACG<u>C</u>AAACGTGGATAAA.

Following the PCR, 20U of DpnI (NEB) was then added directly to the reaction and incubated at 37°C for 1h. A portion of that reaction was then transformed into AG1111 and screened by sequencing. The resulting plasmid was then transformed into *B. subtilis* and subsequently confirmed by antibiotic counter selection and threonine auxotrophy.

### Protein purification

All plasmids encoding genes used for purification were mini-prepped from storage *E. coli* strain AG1111 and freshly transformed into C41(DE3) cells (2) and consequently used for expression of protein (no frozen stocks were used). Briefly, 1L of LB medium was inoculated 1:100 with overnight culture from a single colony. Cells were grown at 37°C with the exception of ClpP and ClpX, which were grown at room temperature. When A600 ∼ 0.6, cells were induced with isopropyl 1-thio-β-D-galactopyranoside to a final concentration of 1mM. Cells were grown for an additional 4-8 h and then cells were harvested by centrifugation, and cell pellets were stored at −80°C. ClpP and ClpX were purified using the IMPACT System (NEB) as described previously (3).

Spx was also purified as previously described (3) with the following modifications: Spx-His was purified from a Ni-NTA column with a 50-500mM imidazole gradient and peak fractions pooled. An N-terminal 6X-His tag was then removed by cleavage with AcTev protease (Life Technologies). Spx was further purified using size exclusion chromatography over an S-300 column in buffer containing 50mM Tris pH 7.5, 50mM KCl, and 10% glycerol. Peak fractions were collected, pooled, and concentrated using an Amicon-10KDa centrifugation column, flash frozen on liquid nitrogen, and stored at −80°C.

His-UgtP in pET28a(+) (PL3521) was purified as follows: Starting from frozen pellets, cells were thawed on ice and re-suspended in Buffer A (50mM Tris pH 8.0, 500mM NaCl, 10mM Imidazole, 10% glycerol). Pefabloc-SC (Sigma) was then added as a protease inhibitor. Re-suspended cells were then lysed by three times by French press at a pressure of 10,000 psi. The lysed cells were then cleared by centrifugation, spinning at 120,000xg for 45 minutes at 4°C. The resulting supernatant was brought up to a volume of 50mL and loaded onto a DynaLoop connected to a DuoFlow F10 FPLC system (Bio-Rad). The supernatant was then applied over two 5mL Bio-Scale Mini Profinity IMAC cartridges connected in series (Bio-Rad). The columns were then washed with 10 column volumes of Buffer A followed by 5 column volumes of Buffer B (50mM Tris pH 8.0, 500mM NaCl, 20mM Imidazole, 10% glycerol). Protein was then eluted off of the columns with 5 column volumes of Buffer C (50mM Tris pH 8.0, 500mM NaCl, 500mM Imidazole, 10% glycerol). Peak fractions were collected and concentrated to a volume of ∼ 1mL using an Amicon-10KDa centrifugation column. The concentrated protein was loaded onto a 1mL loop and applied over a HiPrep 16/60 Sephacryl S-300 HR size exclusion column (GE Healthcare). The column was washed and protein was then eluted off the column in Buffer D (50mM HEPES pH 7.5, 100mM KCl, 10% glycerol). Peak fractions were checked by SDS-PAGE and then aliquot, flash frozen on liquid nitrogen and stored at −80°C.

### UDP-glucose measurements per cell measurements

B. subtilis strains were grown to mid-log, pelleted, and extracted for UDP-glucose. GDP-glucose was added as an internal standard and used to normalize for the extraction efficiency and for quantifications. The samples were run using a LC-MS/MS using a 4000QTRAP. A standard dilution of UDP-glucose was run alongside in 3 replicates to quantify the UDP-glucose in the samples. The average nmole/g was calculated, then applied to cell counts to derive the number of UDP-glucose per cell.

**Additional File 10:**
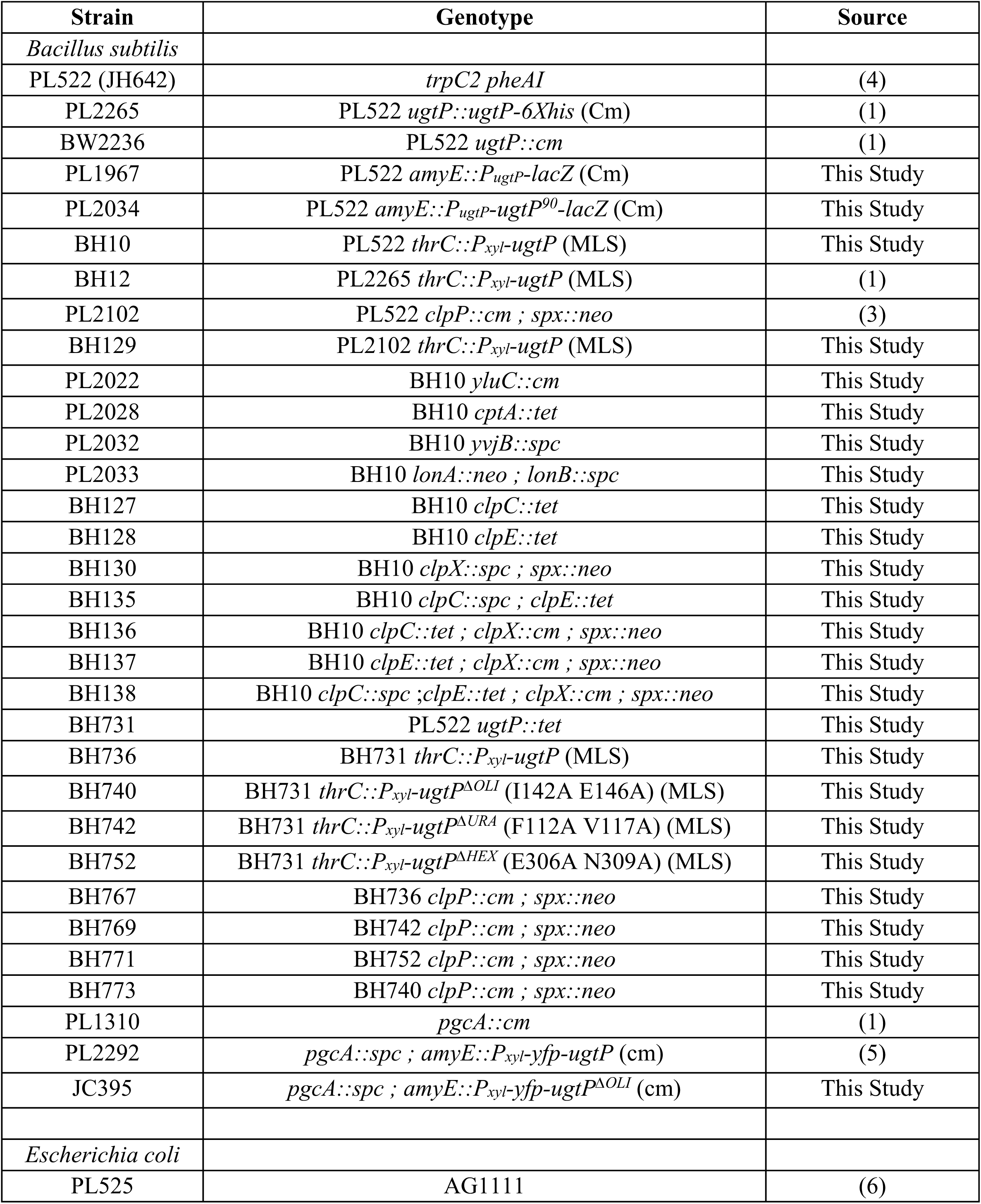
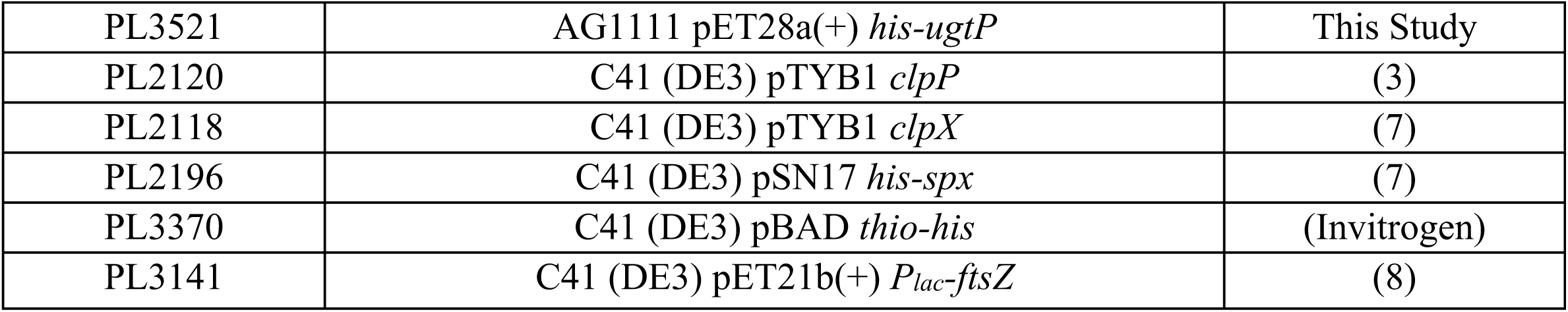
Bacterial Strains Used in This Study.

**Additional File 11:**
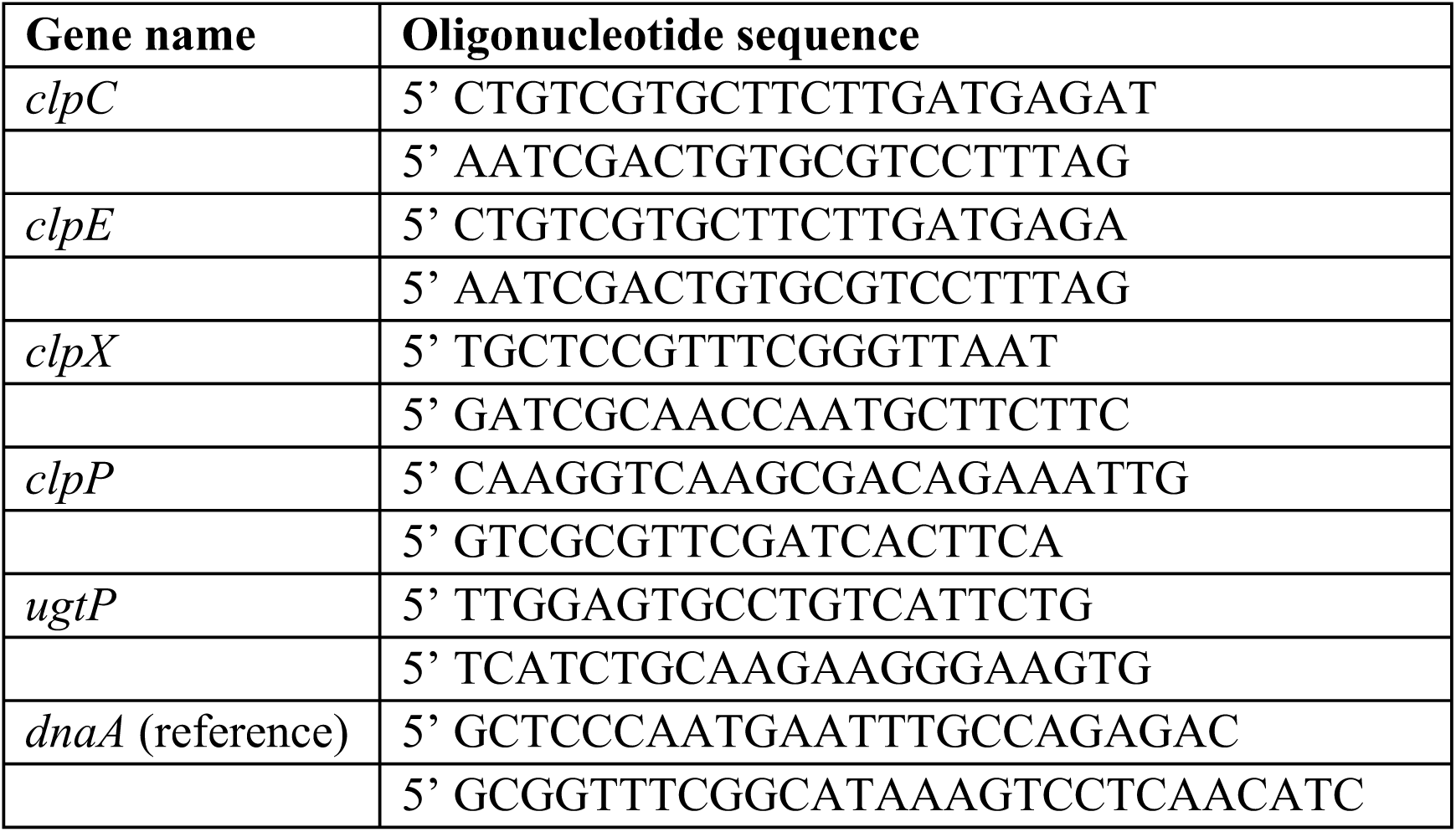
Oligonucleotide Sequences Used for qRT-PCR.

## REFERENCES

1. Schaechter M, Maaloe O, Kjeldgaard NO. 1958. Dependency on medium and temperature of cell size and chemical composition during balanced growth of *Salmonella typhimurium*. J Gen Microbiol 19:592–606.

2. Sargent MG. 1975. Control of cell length in *Bacillus subtilis*. J Bacteriol 123:7–19.

3. Pierucci O. 1978. Dimensions of *Escherichia coli* at various growth rates: model for envelope growth. J Bacteriol 135:559–574.

4. Weart RB, Lee AH, Chien AC, Haeusser DP, Hill NS, Levin PA. 2007. A metabolic sensor governing cell size in bacteria. Cell 130:335–347.

5. Hill NS, Buske PJ, Shi Y, Levin PA. 2013. A moonlighting enzyme links *Escherichia coli* cell size with central metabolism. PLoS Genet 9:e1003663.

6. Klumpp S, Hwa T. 2014. Bacterial growth: global effects on gene expression, growth feedback and proteome partition. Curr Opin Biotechnol 28:96–102.

7. Scott M, Gunderson CW, Mateescu EM, Zhang Z, Hwa T. 2010. Interdependence of cell growth and gene expression: origins and consequences. Science 330:1099–1102.

8. Gerosa L, Kochanowski K, Heinemann M, Sauer U. 2013. Dissecting specific and global transcriptional regulation of bacterial gene expression. Mol Syst Biol 9:658.

9. Klumpp S, Zhang Z, Hwa T. 2009. Growth rate-dependent global effects on gene expression in bacteria. Cell 139:1366–1375.

10. Pedersen S, Bloch PL, Reeh S, Neidhardt FC. 1978. Patterns of protein synthesis in *E. coli*: a catalog of the amount of 140 individual proteins at different growth rates. Cell 14:179–190.

11. Ehrenberg M, Bremer H, Dennis PP. 2013. Medium-dependent control of the bacterial growth rate. Biochimie 95:643–658.

12. Dalebroux ZD, Swanson MS. 2012. ppGpp: magic beyond RNA polymerase. Nat Rev Microbiol 10:203–212.

13. Turgay K, Hahn J, Burghoorn J, Dubnau D. 1998. Competence in *Bacillus subtilis* is controlled by regulated proteolysis of a transcription factor. EMBO J 17:6730–6738.

14. Pan Q, Garsin DA, Losick R. 2001. Self-reinforcing activation of a cell-specific transcription factor by proteolysis of an anti-sigma factor in *B. subtilis*. Mol Cell 8:873–883.

15. Battesti A, Majdalani N, Gottesman S. 2011. The RpoS-mediated general stress response in *Escherichia coli*. Annu Rev Microbiol 65:189–213.

16. Price KD, Roels S, Losick R. 1997. A *Bacillus subtilis* gene encoding a protein similar to nucleotide sugar transferases influences cell shape and viability. J Bacteriol 179:4959–4961.

17. Kennedy EP, Rumley MK. 1988. Osmotic regulation of biosynthesis of membrane-derived oligosaccharides in *Escherichia coli*. J Bacteriol 170:2457–2461.

18. Jorasch P, Wolter FP, Zahringer U, Heinz E. 1998. A UDP glucosyltransferase from *Bacillus subtilis* successively transfers up to four glucose residues to 1,2-diacylglycerol: expression of *ypfP* in *Escherichia coli* and structural analysis of its reaction products. Mol Microbiol 29:419–430.

19. Therisod H, Weissborn AC, Kennedy EP. 1986. An essential function for acyl carrier protein in the biosynthesis of membrane-derived oligosaccharides of *Escherichia coli*. Proc Natl Acad Sci U S A 83:7236–7240.

20. Percy MG, Grundling A. 2014. Lipoteichoic acid synthesis and function in gram-positive bacteria. Annu Rev Microbiol 68:81–100.

21. Reichmann NT, Picarra Cassona C, Monteiro JM, Bottomley AL, Corrigan RM, Foster SJ, Pinho MG, Grundling A. 2014. Differential localization of LTA synthesis proteins and their interaction with the cell division machinery in *Staphylococcus aureus*. Mol Microbiol 92:273–286.

22. Matias VR, Beveridge TJ. 2008. Lipoteichoic acid is a major component of the *Bacillus subtilis* periplasm. J Bacteriol 190:7414–7418.

23. Lazarevic V, Soldo B, Medico N, Pooley H, Bron S, Karamata D. 2005. *Bacillus subtilis* alpha-phosphoglucomutase is required for normal cell morphology and biofilm formation. Appl Environ Microbiol 71:39–45.

24. Chien AC, Zareh SK, Wang YM, Levin PA. 2012. Changes in the oligomerization potential of the division inhibitor UgtP co-ordinate *Bacillus subtilis* cell size with nutrient availability. Mol Microbiol 86:594–610.

25. Gerth U, Kock H, Kusters I, Michalik S, Switzer RL, Hecker M. 2008. Clp-dependent proteolysis down-regulates central metabolic pathways in glucose-starved *Bacillus subtilis*. J Bacteriol 190:321–331.

26. Baker TA, Sauer RT. 2012. ClpXP, an ATP-powered unfolding and protein-degradation machine. Biochim Biophys Acta 1823:15–28.

27. Kress W, Maglica Z, Weber-Ban E. 2009. Clp chaperone-proteases: structure and function. Res Microbiol 160:618–628.

28. Buczek MS, Cardenas Arevalo AL, Janakiraman A. 2016. ClpXP and ClpAP control the *Escherichia coli* division protein ZapC by proteolysis. Microbiology. 162:909–920.

29. Camberg JL, Hoskins JR, Wickner S. 2009. ClpXP protease degrades the cytoskeletal protein, FtsZ, and modulates FtsZ polymer dynamics. PNAS 106(26):10614–10619.

30. Botte C, Jeanneau C, Snajdrova L, Bastien O, Imberty A, Breton C, Marechal E. 2005. Molecular modeling and site-directed mutagenesis of plant chloroplast monogalactosyldiacylglycerol synthase reveal critical residues for activity. J Biol Chem 280:34691–34701.

31. Gerth U, Kirstein J, Mostertz J, Waldminghaus T, Miethke M, Kock H, Hecker M. 2004. Fine-tuning in regulation of Clp protein content in *Bacillus subtilis*. J Bacteriol 186:179–191.

32. Nakano S, Zheng G, Nakano MM, Zuber P. 2002. Multiple pathways of Spx (YjbD) proteolysis in *Bacillus subtilis*. J Bacteriol 184:3664–3670.

33. Zhou Y, Gottesman S, Hoskins JR, Maurizi MR, Wickner S. 2001. The RssB response regulator directly targets sigma(S) for degradation by ClpXP. Genes Dev 15:627–637.

34. Langklotz S, Narberhaus F. 2011. The *Escherichia coli* replication inhibitor CspD is subject to growth-regulated degradation by the Lon protease. Mol Microbiol 80:1313–1325.

35. Schakermann M, Langklotz S, Narberhaus F. 2013. FtsH-mediated coordination of lipopolysaccharide biosynthesis in *Escherichia coli* correlates with the growth rate and the alarmone (p)ppGpp. J Bacteriol 195:1912–1919.

36. Weart RB, Nakano S, Lane BE, Zuber P, Levin PA. 2005. The ClpX chaperone modulates assembly of the tubulin-like protein FtsZ. Mol Microbiol 57:238–249.

37. Sauer RT, Baker TA. 2011. AAA+ proteases: ATP-fueled machines of protein destruction. Annu Rev Biochem 80:587–612.

38. Pan Q, Losick R. 2003. Unique degradation signal for ClpCP in *Bacillus subtilis*. J Bacteriol 185:5275–5278. 25

39. Wiegert T, Schumann W. 2001. SsrA-mediated tagging in *Bacillus subtilis*. J Bacteriol 183:3885–3889.

40. Zhu J, Winans SC. 2001. The quorum-sensing transcriptional regulator TraR requires its cognate signaling ligand for protein folding, protease resistance, and dimerization. Proc Natl Acad Sci U S A 98:1507–1512.

41. Salahudeen AA, Thompson JW, Ruiz JC, Ma HW, Kinch LN, Li Q, Grishin NV, Bruick RK. 2009. An E3 ligase possessing an iron-responsive hemerythrin domain is a regulator of iron homeostasis. Science 326:722–726.

42. Vashisht AA, Zumbrennen KB, Huang X, Powers DN, Durazo A, Sun D, Bhaskaran N, Persson A, Uhlen M, Sangfelt O, Spruck C, Leibold EA, Wohlschlegel JA. 2009. Control of iron homeostasis by an iron-regulated ubiquitin ligase. Science 326:718–721.

43. Hecker M. 2003. A proteomic view of cell physiology of *Bacillus subtilis*--bringing the genome sequence to life. Adv Biochem Eng Biotechnol 83:57–92.

44. Perego M, Spiegelman GB, Hoch JA. 1988. Structure of the gene for the transition state regulator, *abrB*: regulator synthesis is controlled by the *spo0A* sporulation gene in *Bacillus subtilis*. Mol Microbiol 2:689–699.

45. Ireton K, Rudner DZ, Siranosian KJ, Grossman AD. 1993. Integration of multiple developmental signals in *Bacillus subtilis* through the Spo0A transcription factor. Genes Dev 7:283–294.

46. Jaacks KJ, Healy J, Losick R, Grossman AD. 1989. Identification and characterization of genes controlled by the sporulation-regulatory gene *spo0H* in *Bacillus subtilis*. J Bacteriol 171:4121–4129.

47. Hill NS, Kadoya R, Chattoraj DK, Levin PA. 2012. Cell size and the initiation of DNA replication in bacteria. PLoS Genet 8:e1002549.

48. Zhang X, Bremer H. 1995. Control of the *Escherichia coli rrnB* P1 promoter strength by ppGpp. J Biol Chem 270:11181–11189.

49. Pfaffl MW. 2001. A new mathematical model for relative quantification in real-time RT-PCR. Nucleic Acids Res 29:e45.

50. Levin, P. A. 2002. Light microscopy techniques for bacterial cell biology. In Phillipe Sansonetti and Arturo Zychlinsky (Eds.) Methods in Microbiology 31: Molecular Cellular Microbiology, 115–132. Academic Press Ltd., London

## Fig A12: Supplemental References

1. Weart RB, Lee AH, Chien AC, Haeusser DP, Hill NS, Levin PA. 2007. A metabolic sensor governing cell size in bacteria. Cell 130:335–347.

2. Miroux B, Walker JE. 1996. Over-production of proteins in *Escherichia coli*: mutant hosts that allow synthesis of some membrane proteins and globular proteins at high levels. J Mol Biol 260:289–298.

3. Weart RB, Nakano S, Lane BE, Zuber P, Levin PA. 2005. The ClpX chaperone modulates assembly of the tubulin-like protein FtsZ. Mol Microbiol 57:238–249.

4. Perego M, Spiegelman GB, Hoch JA. 1988. Structure of the gene for the transition state regulator, *abrB*: regulator synthesis is controlled by the *spo0A* sporulation gene in Bacillus subtilis. Mol Microbiol 2:689–699.

5. Chien AC, Zareh SK, Wang YM, Levin PA. 2012. Changes in the oligomerization potential of the division inhibitor UgtP co-ordinate *Bacillus subtilis* cell size with nutrient availability. Mol Microbiol 86:594–610.

6. Ireton K, Rudner DZ, Siranosian KJ, Grossman AD. 1993. Integration of multiple developmental signals in *Bacillus subtilis* through the Spo0A transcription factor. Genes Dev 7:283–294.

7. Nakano S, Nakano MM, Zhang Y, Leelakriangsak M, Zuber P. 2003. A regulatory protein that interferes with activator-stimulated transcription in bacteria. Proc Natl Acad Sci U S A 100:4233–4238.

8. Buske PJ, Levin PA. 2012. Extreme C terminus of bacterial cytoskeletal protein FtsZ plays fundamental role in assembly independent of modulatory proteins. J Biol Chem 287:10945–10957.

